# Comparative Genomic Analysis of *Campylobacter rectus* and Closely Related Species

**DOI:** 10.1101/2024.07.26.605372

**Authors:** Casey Hughes Lago, Dana Blackburn, Meghan Kinder Pavlicek, Deborah S. Threadgill

## Abstract

*Campylobacter rectus* is a gram-negative, anaerobic bacterium strongly associated with periodontitis. It also causes various extraoral infections and is linked to adverse pregnancy outcomes in humans and murine models. *C. rectus* and related oral *Campylobacters* have been termed “emerging *Campylobacter* species” because infections by these organisms are likely underreported. Previously, no comparative methods have been used to analyze more than single *C. rectus* strains and until recently, very few *C. rectus* genomes have been publicly available. More sequenced genomes and comparative analyses are needed to study the genomic features and pathogenicity of this species. We sequenced eight new *C. rectus* strains and used comparative methods to identify regions of interest. An emphasis was put on the type III flagellar secretion system (T3SS), type IV secretion system (T4SS), and type VI secretion system (T6SS) because these protein complexes are important for pathogenesis in other *Campylobacter* species. RAST, BV-BRC, and other bioinformatics tools were used to assemble, annotate, and compare these regions in the genomes. The pan-genome of *C. rectus* consists of 2670 genes with core and accessory genomes of 1429 and 1241 genes, respectively. All isolates analyzed in this study have T3SS and T6SS hallmark proteins, while five of the isolates are missing a T4SS system. Twenty-one prophage clusters were identified across the panel of isolates, including four that appear intact. Overall, significant genomic islands were found, suggesting regions in the genomes that underwent horizontal gene transfer. Additionally, the high frequency of CRISPR arrays and other repetitive elements has led to genome rearrangements across the strains, including in areas adjacent to secretion system gene clusters. This study describes the substantial diversity present among *C. rectus* isolates and highlights tools/assays that have been developed to permit functional genomic studies. Additionally, we have expanded the studies on *C. showae* T4SS since we have two new *C. showae* genomes to report. We also demonstrate that unlike *C. rectus*, *C showae* does not demonstrate evidence of intact T6SS except for the strain CAM. The only strain of sequenced *C. massilensis* has neither T4SS or T6SS.

## 1 INTRODUCTION

*Campylobacter rectus* (formerly known as *Wolinella recta*) is a gram-negative, rod-shaped bacterium discovered in 1981 (Anne. C. R. Tanner et al., 1981). It is anaerobic and requires formate and fumarate for growth in liquid media. This organism is an oral pathogen strongly associated with periodontitis and has been identified in 90% of adults with periodontitis, compared with 10% of healthy individuals and 20% of individuals with gingivitis (Macuchi & Tanner, 1999).

In addition to being found in the mouth, *C. rectus* also causes extraoral infections. In recent years there have been case reports of severe and invasive infection caused by *C. rectus,* including some that led to fatalities. The organism has been associated with Barrett’s esophagus, gastroenteritis, appendicitis, Crohn’s disease, ulcerative colitis, empyema thoracis, cerebral microbleeds in stroke patients, peritonitis, and abscesses in the bone (Lam et al., 2011)(Veyrine et al., 2019)(Shiga et al., 2020). This pathogen has also been shown to cause Lemierre’s syndrome, which occurs when a bacterial throat infection becomes systemic and results in sepsis (Hung, Teng, Lau, & Woo, 2019). *C. rectus* is also found in the canine oral cavity and led to pyogenic extensor tenosynovitis in a human patient after a dog bite (Kato Yukio et al., 2011) (Doub, 2021). Additionally, *C. rectus* has also been linked to adverse pregnancy outcomes, including low birth weight and pre-term labor in humans and fetal growth restriction in murine models (Yeo Alvin et al., 2005) (Bobetsis Yiorgos, Barros, & Offenbacher, 2006).

Much research has been completed to study the pathogenesis of *C. jejuni*, *C. coli,* and other enteric *Campylobacter* species. However, less is known about the non-jejuni/coli (njc)- *Campylobacters* including *C. rectus* and related oral organisms such as *C. showae, C. concisus, C. gracilis, C. curvus,* and *C. ureolyticus*. These organisms have been termed “*emerging Campylobacter species*”, which cause infections that are likely to be underreported (Costa & Iraola, 2019). Other njc*-Campylobacters* including *C. showae* and *C. concisus* were also primarily associated with periodontitis like *C. rectus* but have recently been associated with gastroenteritis, irritable bowel disease (IBD), and cancer (Kaakoush, Castaño-Rodríguez, Man, & Mitchell, 2015) (Castaño-Rodríguez, Kaakoush, Lee, & Mitchell, 2017) (Warren et al., 2013).

When this study was initiated, there was a dearth of publicly available *C rectus* genomes. In particular, one ATCC strain known as 33238 and renamed as RM3267 was present through the HOMD website (https://www.homd.org). There are currently 14 publicly available genomes published for *C. rectus* including strains from human, canine, and feline hosts (Table 1). More recently, several unique additional strains have been added, but two are duplications of ATCC 33238 and designated as FDAARGOS_1549. However, no comparative methods have been used to analyze more than single *C. rectus* strains. This lack of genomic information prevents exploration of intraspecific genetic variability and evolution and limits our ability to study pathogenesis.

**Table 1.**
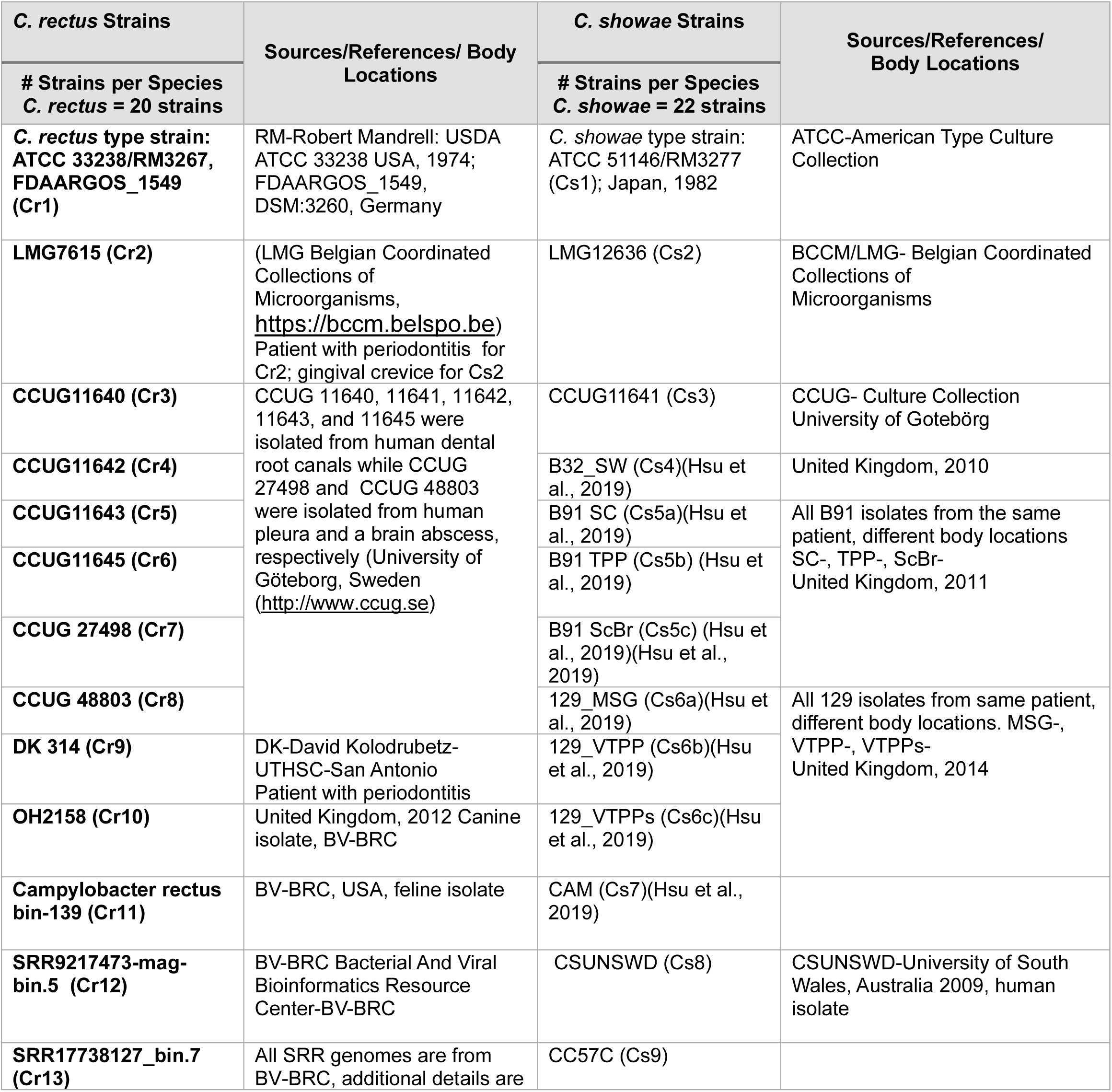
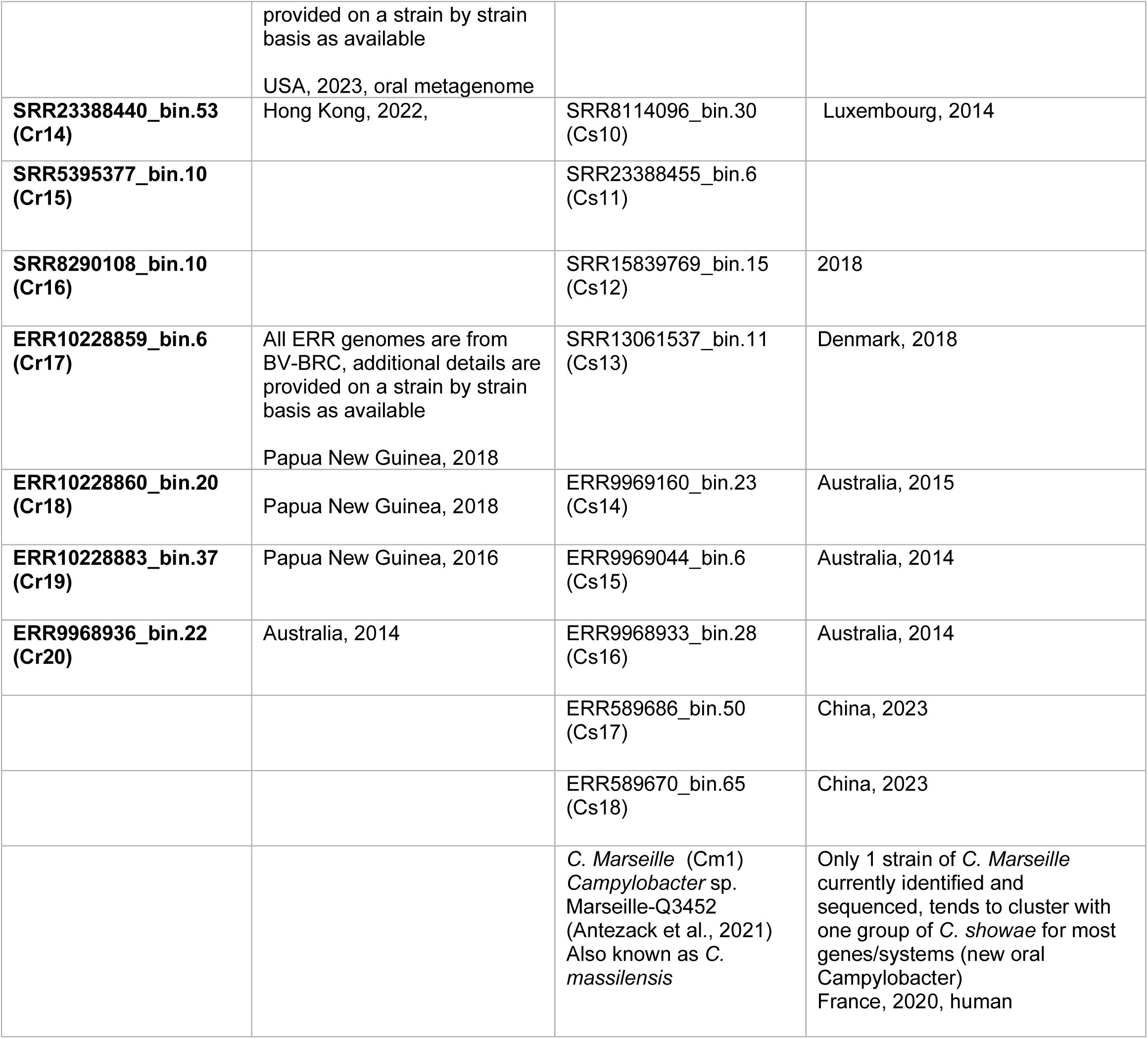
List of current publicly available *C. rectus*, *C. showae and C. massilensis* isolates.

Recently, a comparative and functional genomic analysis was performed for 11 *C. showae* isolates that demonstrated only strains containing T4SS and S-layer proteins were able to adhere to and invade colonic epithelial cells (Hsu et al., 2019). In *C. jejuni*, the T3SS is required for full invasion of host cells, and the T6SS was shown to cause erythrocyte cytotoxicity and enhance host cell adherence and invasion (Lertpiriyapong et al., 2012) (Bleumink-Pluym, Van Alphen, Bouwman, Wösten, & Van Putten, 2013). In *C. jejuni* 81-176, a mutation in VirB11, a T4SS gene located on a plasmid, was shown to reduce adherence and invasion of host cells (Bacon et al., 2000)

Through comparative methods and experimental approaches, we outline the pathogenicity and functionality of 12 *C. rectus* genomes (3 unique publicly available genomes and 9 newly sequenced genomes) and 2 *C. showae* genomes (newly sequenced) from a collection of isolates (Table 2). Three publicly available *C rectus* genomes were used within the analysis as that is what was available at the time of the study. The ATCC 33238 strain maintained in our laboratory was also sequenced, since this strain was used for mutant strain creation. An emphasis was put on secretion systems and virulence factors. Analysis was performed using RAST, PATRIC (now BV-BRC), Island Viewer 4, BRIG, and PHASTER to assemble, annotate, compare, and assess genomic islands and prophage regions in the genomes. Within this panel of genomes, we found that all *C. rectus* strains with complete genomes had T3SS and T6SS systems (determined by presence of hallmark genes) while five of the isolates were lacking a T4SS. With the inclusion of additional *C. rectus* strains in BV-BRC, we have a total of 20 complete or partial genomes for *C rectus* (Table 1). Concurrently we investigated two additional *C. showae* genomes versus those already published and/or publicly available in BV-BRC for T4SS and T6SS as well as T1SS. This results in 22 unique genomes for *C. showae*. (Table 1).

**Table 2.** List of *C. rectus* and *C. showae* isolates sequenced in this study and the three public genomes utilized. *Publicly available genomes.

| STRAIN/Isolate | Contigs | Genome Status | GC Content | Genome Length | No of Genes | Location Isolated |
| --- | --- | --- | --- | --- | --- | --- |
| *C. rectus* CCUG 11645 | 62 | WGS | 44.784 | 2519872 | 3110 | Human dental root canal, apical periodontitis |
| *C. rectus* CCUG 11643 | 62 | WGS | 44.798 | 2532559 | 3123 | Human dental root canal, apical periodontitis |
| *C. rectus* CCUG 11642 | 36 | WGS | 44.901 | 2458336 | 2995 | Human dental root canal |
| *C. rectus* CCUG 11640 | 46 | WGS | 44.819 | 2498484 | 3094 | Human dental root canal, apical periodontitis |
| *C. rectus* CCUG 48803 | 51 | WGS | 44.796 | 2559426 | 3158 | Human cerebrospinal fluid, brain abscess |
| *C. rectus* ATCC 33238 | 73 | WGS | 44.775 | 2546747 | 3104 | Human periodontal pocket |
| *C. rectus* 314 | 55 | WGS | 44.878 | 2523211 | 3136 | Isolate from clinic |
| *C. rectus* LMG 7615 | 48 | WGS | 44.766 | 2544523 | 3099 | Human periodontal pocket |
| *C. rectus* CCUG 27498 | 46 | WGS | 45.138 | 2397792 | 2971 | Human pleura |
| *C. rectus* OH2158* | 91 | WGS | 44.928 | 2491340 | 3085 | Canine oral cavity |
| *C. rectus* SRR9217473-mag-bin.5* | 263 | WGS | 45.907 | 2020017 | 2587 | Human oral cavity |
| *C. rectus* ATCC 33238* | 1 | Complete | 44.71 | 2571679 | 3081 | Human oral cavity |
| *C. showae* 12636 | 27 | WGS | 45.61 | 2287115 | 2469 | Human gingival crevice |
| *C. showae* 11641 | 31 | WGS | 45.831 | 2146162 | 2374 | Human dental root canal |

## 2 MATERIALS AND METHODS

### Bacterial Strains and Culture Conditions

See Table 2 for a complete list of isolates used within the study and related information. All strains were grown in mycoplasma formate fumarate (MFF) media (Mycoplasma Broth Base 20 g/L, 0.2% sodium formate, 0.2% ammonium fumarate) or brain heart infusion (BHI) formate fumarate media (Brain Heart Infusion 38.75 g/L, 0.2% sodium formate, 0.2% ammonium fumarate). Strains were incubated at 37°C in static anaerobic (85% N_2_, 10% H_2_, 5% CO_2_) conditions. The two *C. showae* strains were also grown as described.

### Sequencing, Assembly, and Annotation of Genomes

All isolates were sequenced by the Texas A&M Institute for Genome Sciences and Society Genomics Core Facility. Purified genomic DNA samples were quantified using the Qubit 2.0 High Sensitivity dsDNA assay (LifeTechnologies) and diluted to a starting concentration of 15-20 ng/µL (210-280 ng). Sequencing libraries were prepared using the NEXTflex Rapid DNA-Seq Kit (Bio Scientific) following the manufacturers protocol. Briefly, DNA samples were first fragmented with NEXTflex Enzymatic DNA Fragmentation Enzyme and buffer. Fragmented DNA was repaired, adenylated, and then ligated to adapters (25 uM NEXTflex-HT Barcodes). After Ampure XP bead cleanup, libraries were amplified with 5 cycles of PCR (30 sec at 98°C, 30 sec at 65°C, and 60 sec at 72°C). Library size was assessed on a D1000 ScreenTape on the Agilent TapeStation with an average DNA peak at ∼425 bp. Samples were quantified using the Qubit 2.0 High Sensitivity dsDNA assay (LifeTechnologies), normalized to 4nM, then pooled for sequencing. Sequencing was performed on an Illumina MiSeq with a 2×150 cycle version 2 Micro Reagent Kit. Sequencing data was then uploaded to the Illumina cloud service, BaseSpace, for storage and analysis. A total of 4.6 million reads were generated with each sample accounting for between 874,000-1,300,000 reads. Each sample was analyzed using SPAdes Genome Assembler version 3.9.0 to generate contig fasta files. The genomes were annotated in the Bacterial and Viral Bioinformatics Resource Center (BV-BRC) using the Rapid Annotation using Subsystem Technology (RASTk) pipeline (https://rast.nmpdr.org) (Brettin et al., 2015; Wattam et al., 2017).

### BRIG, Phylogeny, Pangenome, and Core Protein Identification

To create a graphical representation of similarities between the genomes, the BLAST Ring Image Generator (BRIG) was used (Alikhan, Petty, Ben Zakour, & Beatson, 2011). *C. rectus* 48803 was used as a reference since it was the largest assembly. A pipeline in BV-BRC was used to build a phylogenetic tree based on 1000 genes. This pipeline uses amino acid and nucleotide sequences from a defined number of BV-BRC’s global protein families (PGFams), which are picked randomly, to build an alignment, and then generate a tree based on the differences within those selected sequences. *C. showae* CCUG 11641 and LMG 12636 were used as an outgroup and FigTree was used to generate the final image (Rambout, 2007). The Orthologous Average Nucleotide Identity Tool (OAT) was utilized to perform pairwise comparisons (Lee, Ouk Kim, Park, & Chun, 2016)

The pan-genome of the panel studied was determined using BV-BRC’s Protein Family Sorter tool. Core proteins of interest and proteins related to secretion systems were identified by searching the genomes using the “features” tab.

### Identification of Prophage, Genomic Islands, and Plasmids

Isolates were analyzed using PHASTER (http://www.phaster.ca) to identify both complete and incomplete prophage regions in the genomes (Arndt et al., 2016). Genomic islands (GI) were predicted by IslandViewer 4 (http://www.pathogenomics.sfu.ca/islandviewer/upload/), which is an integrated interface for computational identification and visualization of genomic islands that utilizes four different prediction methods: IslandPath-DIMOB, SIGI-HMM, IslandPick and Islander (Bertelli et al., 2017).

To determine if any isolates contained plasmids, pulse field gel electrophoresis (PFGE) was performed using the CHEF Bacterial Genomic DNA Plug Kit (BioRad-Laboratories). Briefly, each strain was grown anaerobically as previously described in 7 mL BHI-FF liquid media for 72 h. The OD_600_ was adjusted for each sample to 0.8 and cultures were centrifuged at room temperature (10 min, 3,500 rpm). Supernatants were removed and pellets were resuspended in 100 µL of cell suspension buffer prior to adding 100 µL prewarmed 2% CleanCut agarose and plugs were then treated with lysozyme, proteinase K, and washed according to the manufacturer’s instructions. Once ready, the plug slices were loaded into the appropriate wells of a 1% agarose gel (Pulsed Field Certified Agarose, BioRad-Laboratories). The gel was electrophoresed for 18 h in 0.5× TBE at an angle of 120° with initial and final switch times of 2 and 10 seconds respectively. After the electrophoresis run was complete, the gel was stained in 225 mL of 0.5× TBE with 60 µL of Gel Red (10,000X) and band pattern was observed under UV illumination.

### Development of Tools for Functional Genomic Studies of *C. rectus* secretion systems

Although not a primary focus of the comparative genomics detailed in this study, targeting vectors were created to knock-out a single gene for each secretion system. The design of these targeting vectors will be detailed in later publications focusing on mutant phenotype characterizations. Mutations were made in the ATCC 33238 strain initially since this was the first genomic isolate for which entire genome sequence data was available. All mutations were complete gene deletions, and include *ciaB* (T3SS), *virB9* (T4SS) and *hcp* (T6SS) (Supplemental Table 6). The *ciaB* deletion mutant has been the most fully examined and is described further in (Hughes Lago et al., Manuscript in Preparation). Another tool that was developed for *C. rectus* is real-time PCR including establishing a set of genes that could serve as internal controls for mutant strain analysis.

### RT qPCR

See supplemental methods for RNA isolation, cDNA synthesis, and DNase treatment. To evaluate candidate reference genes, four biological replicates per media type were utilized. To test gene expression, three biological replicates per strain were used. Both ATCC 33238 and strain 314 were used for the reference gene studies. To determine Cq values for all genes, a Bio-Rad C1000 Touch Thermal Cycler with the CFX96 Real-Time System (Bio-Rad Laboratories) was used. The iTaq Universal SYBR Green One-Step Kit (Bio-Rad Laboratories) was used according to manufacturer instructions for 10 mL reactions using 200 ng of RNA from each sample. The expression patterns of the following candidate reference genes were examined: 16S ribosomal RNA (*16S*), DNA gyrase (*gyrA*), recombinase A (*recA*), serine hydroxymethyltransferase (*glyA*), RNA polymerase (*rpoA*), and RNA polymerase sigma factor (*rpoD*); commonly used as validated reference genes in other bacteria (Rocha, Santos, & Pacheco, 2015). Reactions were completed with primers for the six reference genes (Supplemental Table 1) with no template controls included for all primer sets. Additionally, a no reverse transcriptase control was included for all samples using the *glyA* (33238) and *rpoA* (314) primer sets. For the thermal cycler, the following parameters were used: 10 min at 50°C, 1 m at 95°C, followed by 40 cycles of 95°C 15 s, 55°C 15 s, 72°C 30 s, with a 65-95°C melt curve analysis with 0.5°C increase and 5 s per cycle.

### Phenotypic Assessments

Biofilm formation was examined for all *C. rectus* strains during stationery culture in anaerobic conditions using 96-well, flat-bottom, polystyrene microplates (Costar) and crystal violet stain, as previously outlined (O’Toole, 2011). *Pseudomonas aeruginosa* PAK was used as a positive control. DNA uptake was another phenotype explored for all *C. rectus* strains. See supplemental methods for detailed protocol.

## 3 RESULTS AND DISCUSSION

### Sequencing and Assembly

Nine *C. rectus* and two *C. showae* strains were sequenced using Illumina MiSeq. All sequenced genomes were estimated to be >99% complete according to BV-BRC genome summary reports with course consistency 97.2% or greater, and fine consistency 96.4% or greater for all sequenced isolates (Table 2). Each assembly had 73 contigs or less. N50 scores ranged from 9071 to 368,575. GC content ranged between 44.78 to 45.91%. The two publicly available genomes analyzed had the lowest contiguity with SRR9217473-mag-bin.5 and OH2158 having 263 and 91 contigs respectively.

### BRIG, Phylogeny, Pangenome, Prophage, and Genomic Islands

The *C. rectus* isolates were compared using BRIG and a phylogenetic tree (Figures 1 and 2). Within the phylogeny, all isolates clustered into two distinct clades. The first containing 11645, 11643, 11642, 11640, 48803, and 314, and the second containing 33238, 7615, OH2158, 27489, and SRR921743. Both 33238 and 7615 grouped together, appearing highly similar. Multiple prophage were found in all the isolates with both complete and incomplete fragments (Table 3). Five complete prophage were identified including four *Pseudomonas* phage PPpW-3 (NC_023006), and one *Escherichia* phage vB_EcoM_ECO1230-10 (NC_027995). CCUG 48803 had the most phage with 1 complete phage, and 3 incomplete phage fragments. The presence of phage contributes to the genetic diversity among these isolates. Multiple genomic islands were also identified indicating regions in the genomes may have undergone horizontal gene transfer from transformation, conjugation, or transduction. *C. rectus* ATCC 33238 has the largest portion of genomic DNA from genomic islands at 10.8% followed by LMG 7615 with 9.65%. *C. rectus* 27498 has the lowest amount of genomic DNA from genomic islands at 2.85% (Table 4).

**Figure 1.**
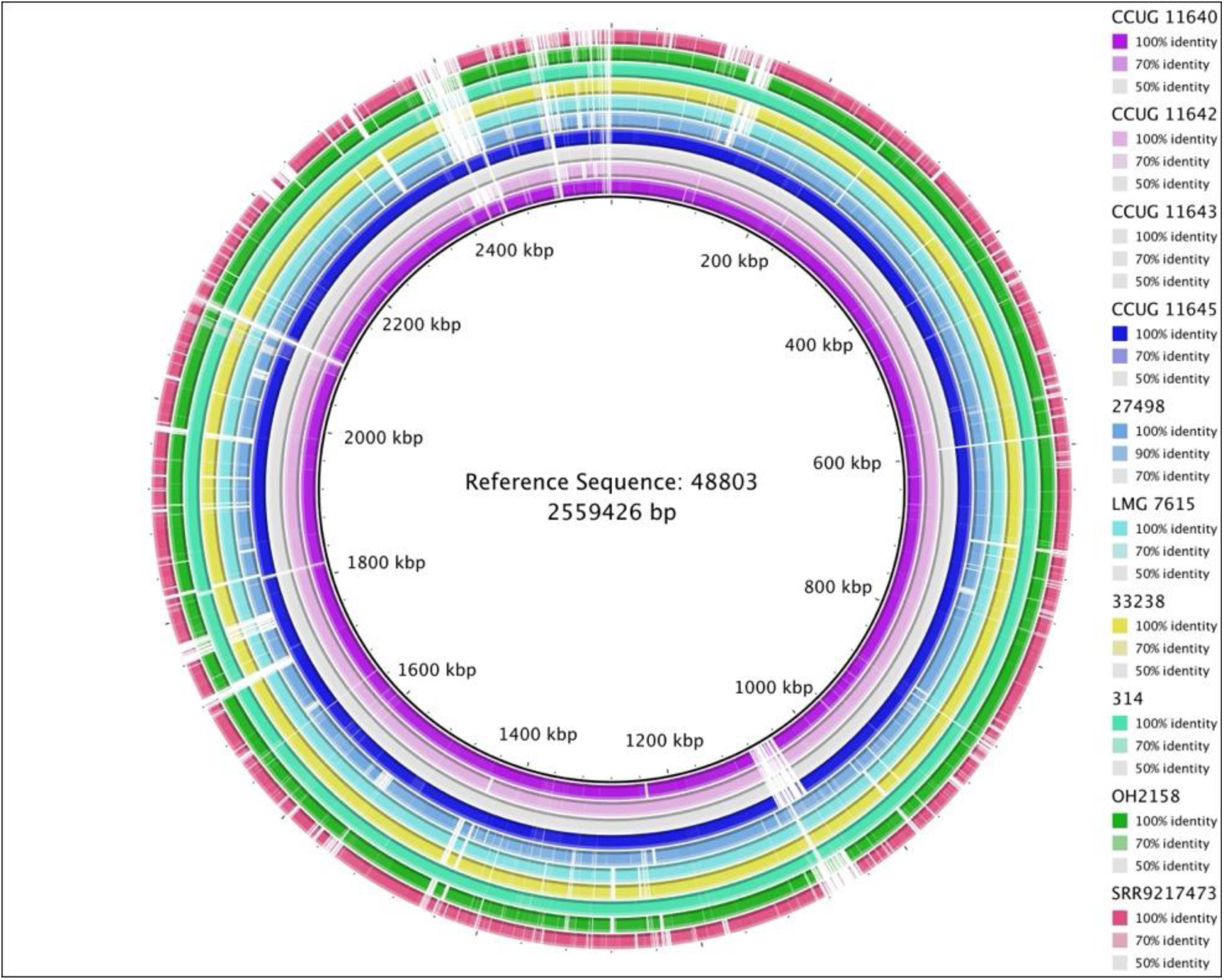
BRIG (Blast Ring Image Generator) showing eight newly characterized *C. rectus* strains aligned to the longest genome (48803).

**Figure 2.**
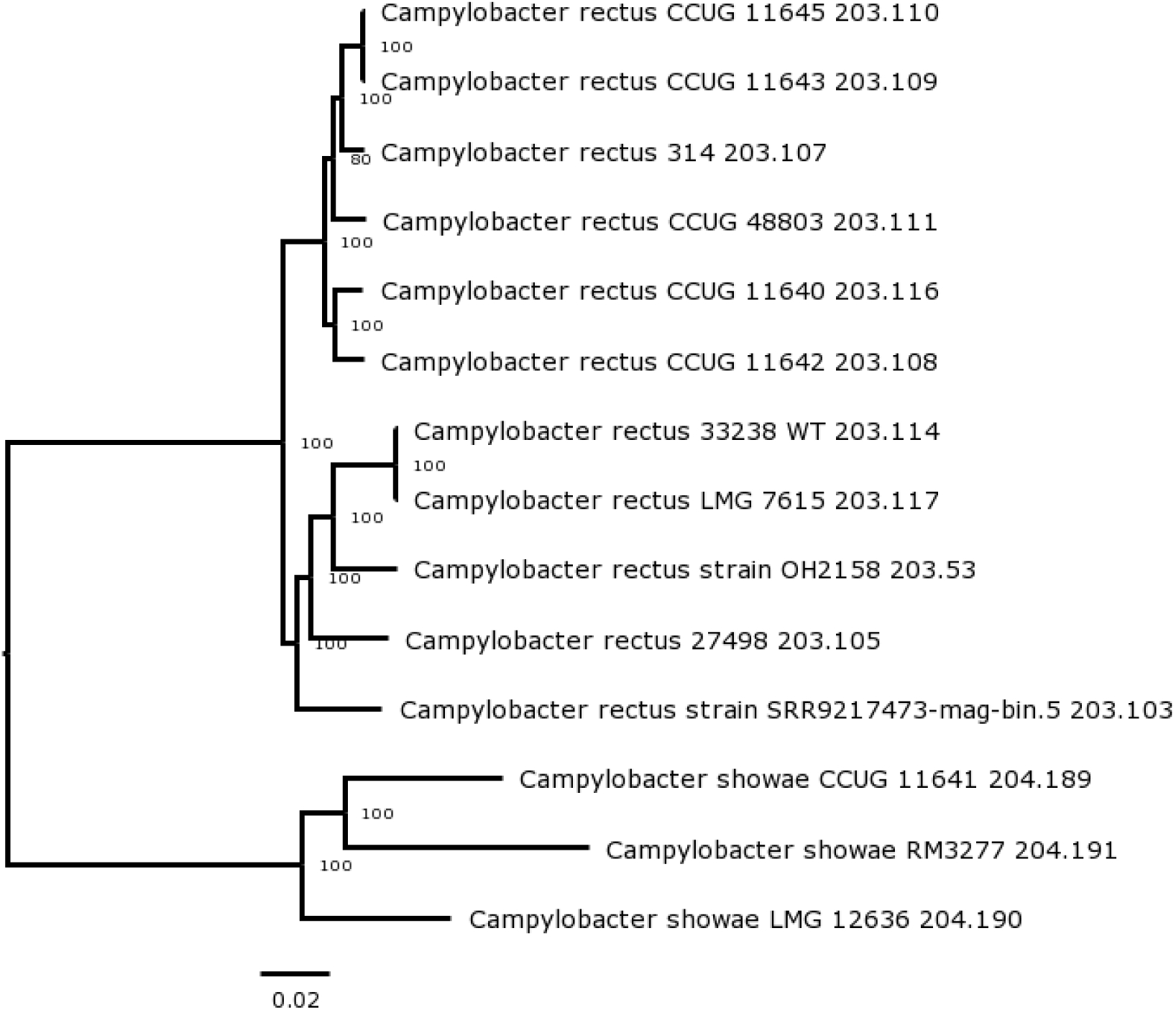
Phylogenetic tree with *C. rectus* and *C. showae* isolates. Bootstrap values are shown at each node.

**Table 3.** Distribution of Genomic Islands (GI) in *C. rectus* isolates.

| STRAIN/Isolate | Size of GI (bp) | Total Length of GI/Genome Size (%) |
| --- | --- | --- |
| *C. rectus* CCUG 11645 | 129295 | 5.1% |
| *C. rectus* CCUG 11643 | 158665 | 6.26% |
| *C. rectus* CCUG 11642 | 145819 | 5.9% |
| *C. rectus* CCUG 11640 | 156764 | 6.27% |
| *C. rectus* CCUG 48803 | 190477 | 7.44% |
| *C. rectus* ATCC 33238 | 274235 | 10.8% |
| *C. rectus* 314 | 174842 | 6.93% |
| *C. rectus* LMG 7615 | 245547 | 9.65% |
| *C. rectus* CCUG 27498 | 68422 | 2.85% |

**Table 4.** Whole and Fragmented Prophage Identified in *C. rectus* Isolates.

|  | Region | Size (Kb) | Completeness | Score | No of Proteins | Position in Genome | Most Common Phage |
| --- | --- | --- | --- | --- | --- | --- | --- |
| *C. rectus*<br>CCUG 11645 | 1 | 15.2 | incomplete | 20 | 8 | 851292-866555 | Cyanophage S-RIM12 (NC_047717) |
|  | 2 | 11.1 | incomplete | 19 | 19 | 2313301-2324420 | Lactobacillus phage phiLdb (NC_022762) |
| *C. rectus*<br>CCUG 11643 | 1 | 15.7 | incomplete | 20 | 8 | 1546549-1562295 | Cyanophage S-RIM12 (NC_047716) |
|  | 2 | 11.1 | incomplete | 30 | 19 | 2359733-2370852 | Mannheimia phage vB_MhS_587AP2 (NC_028743) |
| *C. rectus*<br>CCUG 11642 | 1 | 15.7 | incomplete | 20 | 8 | 16863-32609 | Cyanophage S-RIM12 (NC_047716) |
| *C. rectus*<br>CCUG 11640 | 1 | 15.2 | incomplete | 20 | 8 | 850728-865991 | Cyanophage S-RIM12 (NC_047717) |
|  | 2 | 27.2 | incomplete | 40 | 31 | 1155337-1182561 | Lactobacillus phage phiLdb (NC_022762) |
| *C. rectus*<br>CCUG 48803 | 1 | 15.2 | incomplete | 20 | 8 | 523767-539023 | Cyanophage S-RIM12 (NC_047716) |
|  | 2 | 26.2 | intact | 140 | 42 | 1046511-1072799 | Pseudomonas phage PPpW-3 (NC_023006) |
|  | 3 | 6.3 | incomplete | 40 | 9 | 1077452-1083755 | Geobacillus virus E3 (NC_029073) |
|  | 4 | 11.1 | incomplete | 30 | 18 | 2394871-2406036 | Mannheimia phage vB_MhS_587AP2 (NC_028743) |
| *C. rectus*<br>33238 WT | 1 | 24.5 | intact | 150 | 41 | 83191-107756 | Pseudomonas phage PPpW-3 (NC_023006) |
| *C. rectus*<br>314 | 1 | 36.2 | intact | 150 | 63 | 463372-499651 | Pseudomonas phage PPpW-3 (NC_023006) |
|  | 2 | 15.2 | incomplete | 20 | 8 | 1601400-1616663 | Cyanophage S-RIM12 (NC_047717) |
| *C. rectus*<br>LMG 7615 | 1 | 24.5 | intact | 150 | 42 | 1368107-1392672 | Pseudomonas phage PPpW-3 (NC_023006) |
| *C. rectus*<br>27498 | 1 | 17.6 | incomplete | 40 | 9 | 850297-867984 | Geobacillus virus E3 (NC_029073) |
|  | 2 | 24.9 | intact | 140 | 42 | 859544-884473 | Escherichia phage vB_EcoM_ECO1230-10 (NC_027995) |
|  | 3 | 13.2 | incomplete | 20 | 9 | 1978229-1991480 | Cyanophage S-RIM12 (NC_047716) |
| *C. rectus*<br>OH2158 | 1 | 9 | incomplete | 30 | 10 | 41716-50806 | Phormidium phage MIS-PhV1B (NC_028998) |
|  | 2 | 9.8 | incomplete | 20 | 14 | 15391-725282 | Rhodovulum phage vB_RhKS_P1 (NC_031059) |
| *C. rectus*<br>SRR9217473<br>-mag-bin.5 | 1 | 11 | incomplete | 40 | 13 |  | Synechococcus phage ACG-2014f (NC_026927) |

The pan-genome of the panel studied consists of 2670 genes with core and accessory genomes of 1429 and 1241 genes, respectively. Core proteins identified include DNA Polymerase I and III as well as flagellar genes FlgG, FlhA, FlhB, and FlhF. Fumarate metabolism genes identified include formate dehydrogenase and formate tetrahydrofolate ligase. A large amount of CRISPR arrays and repeat regions were also identified across the isolates (Supplemental Table 4). No significant pairwise comparisons were observed (Supplemental Figure 1)

### Expression Analysis

The 16S ribosomal RNA (*16S*), DNA gyrase (*gyrA*), recombinase A (*recA*), serine hydroxymethyltransferase (*glyA*), RNA polymerase (*rpoA*), and RNA polymerase sigma factor (*rpoD*) genes were chosen as candidate reference genes because they are commonly used as transcriptional reference genes in other bacteria (Rocha, Santos, & Pacheco, 2015). The expression of all candidate genes was examined in 33238 and 314 except for the 16S rRNA gene, where the 16S Cq values were too low to analyze. There was no significant difference in expression levels of the other candidate reference genes between samples grown in MFF and TSBFF, except for *gyrA* levels in 33238; since the difference was minor, *gyrA* was still used in future analyses.

No single reference gene was the most resistant to expression change in different growth media for both strains of *C. rectus* (Supplemental Table 2). However, the genes showing the smallest expression changes in both strains were similar. The top three reference genes for 33238 were *rpoA*, *recA*, and *gyrA*, while the top genes for 314 were *recA*, *rpoD*, and *gyrA* or *rpoA*. Results using NormFinder identified the same top three genes for 33238 as the BestKeeper analysis (Supplemental Table 3). This real-time PCR assay has been used to verify the mutant strains (ATCC 33238 parent strain) discussed in supplemental methods.

### Plasmid Determination and Phenotypic Assessments

Upon completing PFGE, we cannot conclusively report plasmids. Additionally, no embedded plasmids were identified from the genome sequencing. No biofilm formation was observed in any of the strains. DNA uptake experiments did not account for types of DNA uptake (i.e. natural transformation versus conjugation via T4SS) so we cannot make any judgements about this phenotype currently except that it is definitely a varying phenotype across strains (Supplemental Table 5). Further optimized studies with the virB9 mutant and strains lacking T4SS will need to be completed to address the importance of T4SS and Com genes to DNA uptake for *C. rectus*.

### Secretion Systems

In gram-negative bacteria, the type 1 secretion system (T1SS) facilitates secretion across the cell membrane. This process requires ATP hydrolysis and T1SS components broadly including an ABC transporter, periplasmic adaptor protein, and an outer-membrane protein (Spitz et al., 2019). The first T1SS was identified in *E. coli,* which became the classic description of the mechanism. Since then, a variety of T1SS have been characterized and classified into 5 different subgroups (Hodges, Torres, Cunningham, Henderson, & Icke, 2023).

Type 1 secreted products may include bacterial toxins like hemolysins, leukotoxins, proteases, or other proteins including surface-layer proteins (SLP’s). SLP’s allow bacteria to be resistant to complement and antibodies which is a key defense of extracellular pathogens. In *C. fetus*, this system is composed of a SapDEF complex which exhibits homology to the T1SS (Tu, Gaudreau, & Blaser,). T1SS proteins were identified in two different gene clusters in each of the *C. rectus* isolates analyzed (Figure 3 and Supplemental Figure 2). One cluster includes SapC, SapD, SapE, and SapF, flanked by at least one SLP. The other cluster does not have all the Sap genes but does have the structural genes needed for Type 1 secretion namely an ABC transporter, membrane fusion protein (SapE), and LapP*. C. showae* isolates appear to have one or the other (Figure 3 and Supplemental Figure 2). The lack of SLP’s for many *C. showae* strains has been suggested to relate to virulence, since the S-layer expressing strains were assessed as more invasive (Hsu et al., 2019). It should be noted that the T1SS in *C. rectus* containing the SLP does not closely match that of *C. fetus* in that only one SLP (Crs) has been identified rather than several different SapA or B proteins. *C. showae* may be more like *C. fetus* in that several SLPs have been identified for those strains that have S-layer Type I. The SLPs of *C. showae* are substantially smaller than the Crs protein of *C. rectus* (Table 5) and only some of the identified S-layer proteins have homology with the Crs. it will be interesting to investigate the origin of S-layer for *C. showae* since both *C. fetus* and *C. rectus* have this type of structure for all strains and *C. showae* does not.

**Figure 3.**
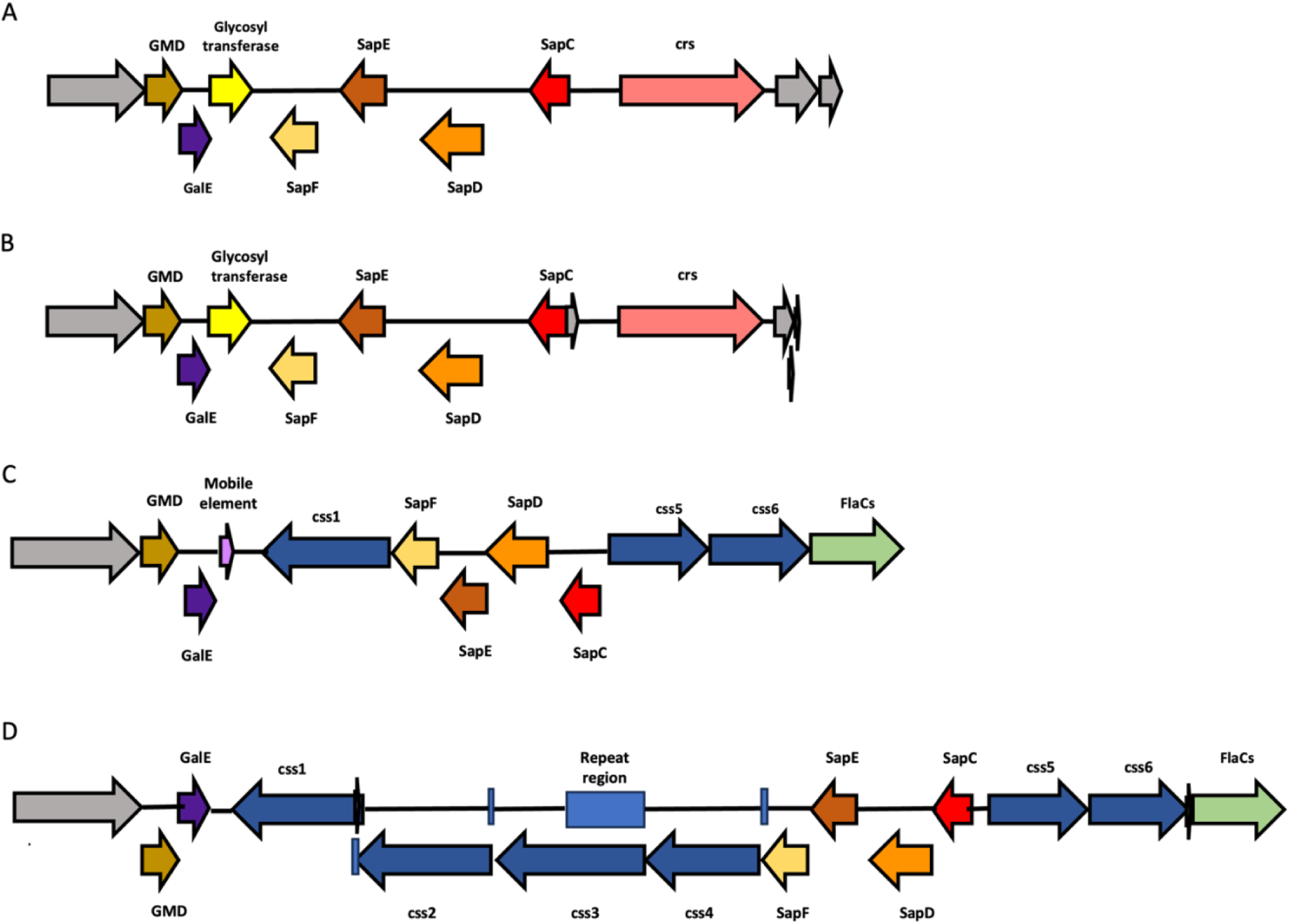
Comparison of T1SS clusters in A) *C. rectus* 7615, B) *C. massilensis,* C) *C. showae* 51145, D) *C. showae* CAM (Gene Abbreviations: GMD: GDP mannose 4,6 dehydratase, GalE: UDP glucose 4 epimerase, SapF: T1S outermembrane protein, TolC Family, efflux transport system, SapE: T1S membrane fusion protein hylD family, SapD: T1 Secretion system ATPase, SapC: peptide transport system permease protein, crs: s-layer protein) Hypothetical proteins are in grey.

**Figure 4.**
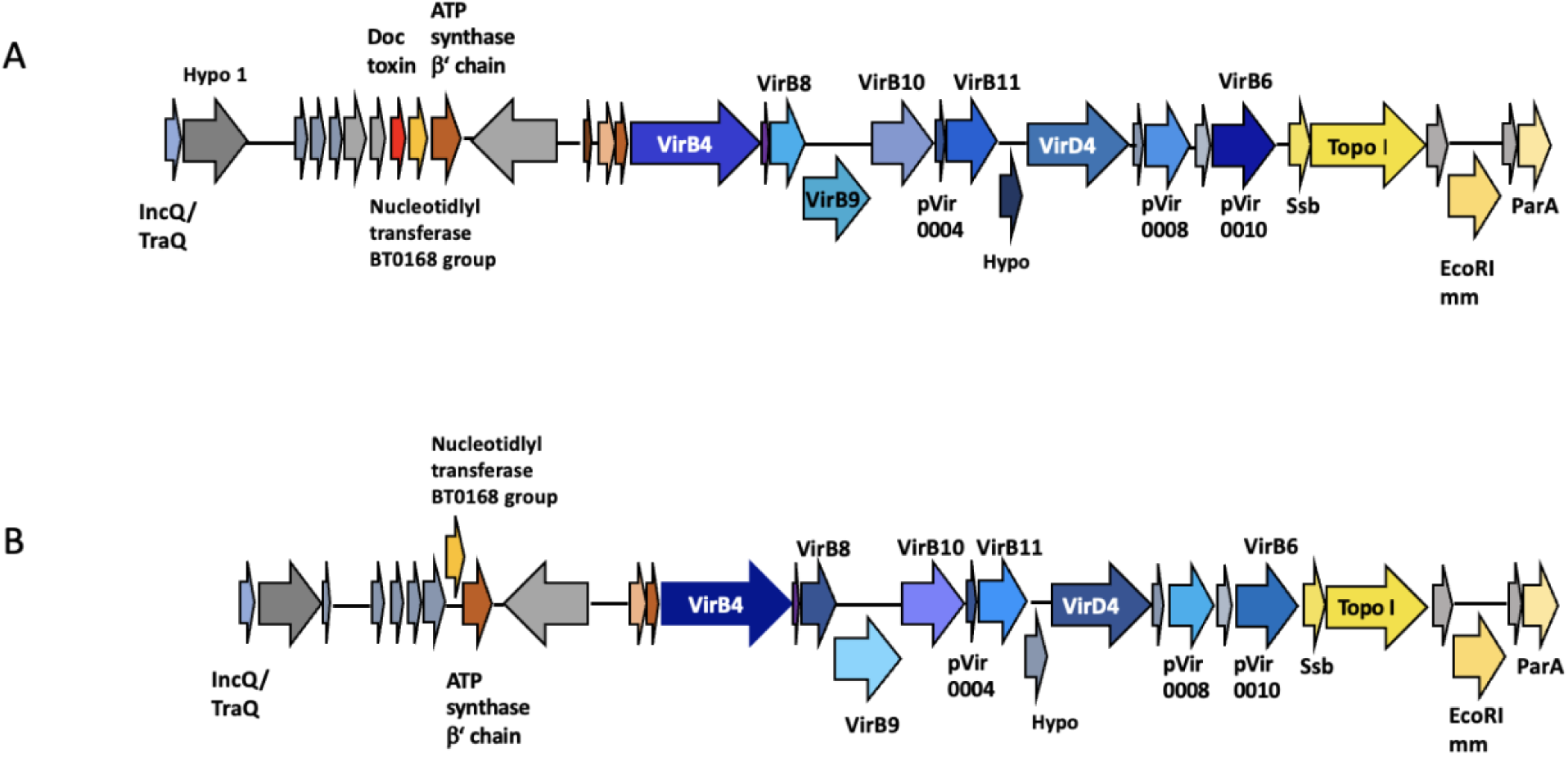
Comparison of T4SS clusters in A) *C. rectus* 7615 B) *C. showae* 11641

**Figure 5.**
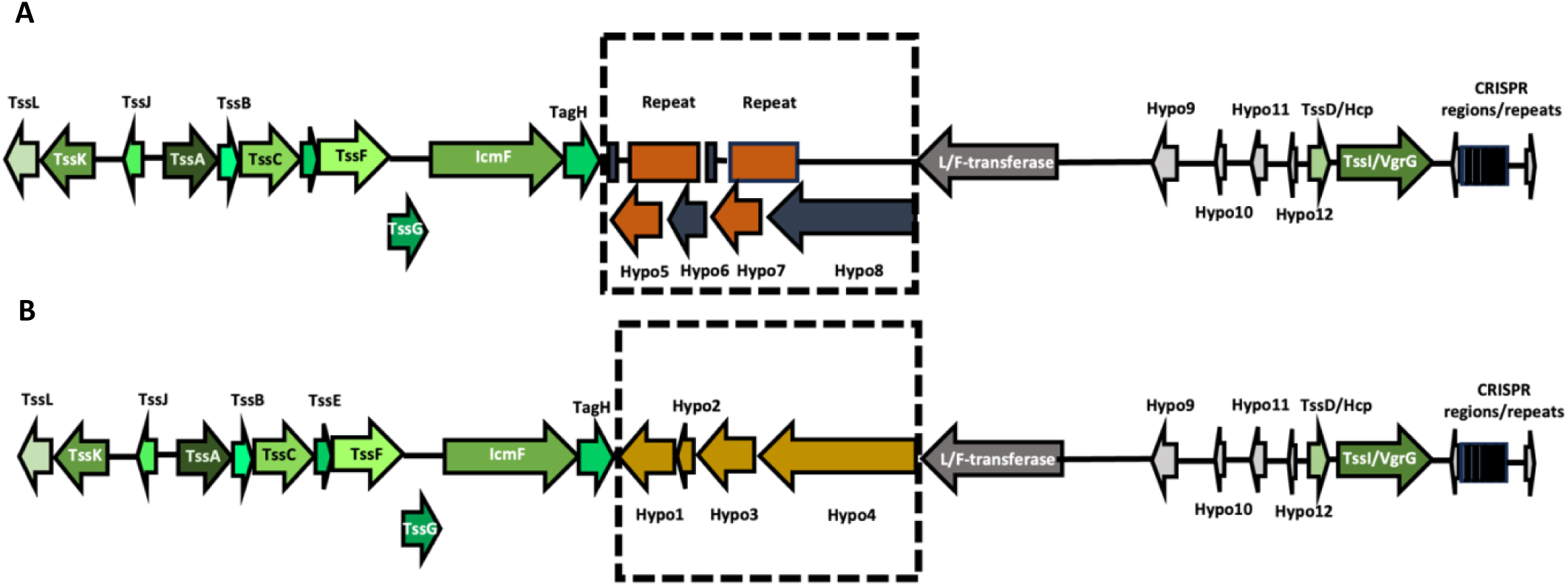
Comparison of T6SS clusters showing different repeat regions in A) *C. rectus* 11643 and B) hypothetical proteins common to both 33238 and CAM (Gene Names and/or Orthologs: TssL; VasF/ImpK, TssK; VasE/ImpJ, TssJ; VasD, TssA; VasJ/ImpA, TssB; ImpB, TssC; VipB/ImpC, TssE; VCA0109, TssF; VasA/ImpG, TssG; VasB, TssM; VasK/IcmF, TagA, L/F-transferase; Leucyl/phenylalanyl-tRNA-- protein transferase, Hypo; hypothetical protein)

**Table 5.** *C. rectus, C. showae* and *C. massilensis* Protein Secretion System Clusters.

| Protein Name/<br>Locus name | Strains / No of strains | Genes found for the<br>secretion system | Unique features |
| --- | --- | --- | --- |
| **Type I Protein Secretion** | All *C. rectus* and *C. showae* strains with complete genomes have this type of protein secretion system, albeit only some *C. showae* have the Type I surface layer secretion system. And only those that lack this system in *C. showae* have the non-Slp secretion system. | *C. massilensis* has both types of T1SS and thus resembles *C. rectus* for these secretion systems | Mentioned because it is the mode for secretion of surface layer proteins, as well as surface layer cytotoxins.<br>Is another Type I system that may impact virulence |
| **Type I Surface Layer Proteins** |  |  |  |
| ***C. rectus* major S-layer protein (crs)** | Cr1-Cr11, Cm1<br>Also complete or partial S-layer clusters in Cr12-18, and Cr20 |  | Full-size crs<br>See figure 3 in main text for visual explanation of the Type I SLP clusters in *C. rectus* and *C. showae*. |
| ***C. showae* major S-layer protein (css)** | Present in Cs1, Cs4, Cs6a-c, Cs7-Cs8 | Since no nomenclature exists currently for S-layer proteins in *C. showae* we propose the css nomenclature for the genes, and Css1-6 for the proteins found. Css1 has been named based on the comparison between strain Cs1 and Cs7 as these have the most similar arrangement of the genes and proteins for *C. showae*. | The two smaller S-layer proteins will be numbered Css5 and Css6 since these are less common than the large proteins, i.e. fewer strains have them. Only Cs1 and Cs7 have both of these smaller S-layer proteins and their Css5 sequences match well as do their Css6 sequences. |
| **SapC, SapD, SapE, SapF** | Cr1-Cr11<br>Cs1, Cs4, Cs6a-c, Cs7-Cs8 | 129_group inclusive | SapC - peptide transport system permease protein<br>SapD – T1 Secretion system ATPase<br>SapE – T1S membrane fusion protein hylD family<br>SapF – T1 Secretion outer membrane protein TolC (efflux transport system, OMF lipoprotein) |
| **Type I non-SLP** | See supplemental figure 2 for the organization of this Type I system |  |  |
| **LapP, LapD, GGDEF and EAL domain proteins,** | Cr1-Cr11, Cm1<br>Cs2-Cs3, Cs5a-5c, Cs9 | B91_group inclusive | LapP- T1SS associated transglutaminase-like cysteine proteinase |
| ABC transporter, Hyl D family member, Agglutination protein |  |  | LapD- membrane bound c-di-GMP receptor<br>In some cases LapD is called GGDEF and EAL domain proteins<br>ABC transporter, transmembrane region: ABC transporter:Peptidase C39, bacteriocin processing<br>Type I secretion membrane fusion protein, HlyD family<br>Agglutination protein |
| T3SS flagellar secretion system | Conserved in flagellated bacteria |  | Related to T3SS Injection needle secretion systems of other Genera of bacteria but only uses flagellar portal apparatus as in *C. jejuni* |
| *Campylobacter* invasion antigen B- CiaB | All strains/species examined appear to have this antigen but this will be described more fully in a CiaB specific paper (C.H. Lago et al., in preparation) |  |  |
| FlaA/B flagellin filament protein | Only one flagellin filament present in the majority of *C. rectus* and *C. showae* strains, and it does not match completely for either FlaA or FlaB of *C. jejuni* and is only annotated as Fla in the genomes<br>Both FlaA and FlaB are longer than the *rectus_showae* flagellins and also the AA differences distinguishing A from B are not uniformly represented in *rectus_showae*. |  |  |
| Fla<br>The flagellin of *C. rectus* | Only one *C. rectus* group for flagellin. | Uniform size for the flagellin subunit |  |
| FlaCs<br>The flagellin of *C. showae* | Two *C. showae* groups of FlaCs known as 129_ group and B91_ group because these related strains are the foundation members of the group | 129_ FlaCs group has a flagellin location close to the Type I S-layer secretion system. 129_ group is Cs1, Cs4, Cs6a-c, Cs7-Cs8<br>B91_ FlaCs group has a flagellin location close to the Pse cluster<br>B91 group is Cs2-Cs3, C5a-5c, Cs9 | See T1SS S-layer Figure for demonstration of this for the 129_group |
| **Pseudaminic Acid Cluster (Pse)**<br>**Involved in flagellin glycosylation and antigenicity of flagellin proteins (Salah Ud-Din & Roujeinikova, 2017)** | PseB, PseC, PseF, PseG, PseH, PseI<br>All *C. rectus* and *C. showae* with flagellin also have the pse cluster somewhere in the genome, suggesting that similar to *C. jejuni* and *H. pylori* Pse is stabilizing for flagellin and required for secretion and assembly of flagella | The motility accessory protein or MAF has been annotated as PseE in *C. showae* 51146 because it is the flagellar glycosyl transferase. Flagellin that fails to be post-translationally modified with Pse fails to assemble into mature flagellin and results in loss of motility. | The Pse cluster is located adjacent to the flagellin in all B91_ FlaCs strains as well in all *C. rectus* strains but is located near LPS related genes and the S-layer proteins in all 129_FlaCs strains |
| **FlaC**<br>**FlaC(s)** | All *C. rectus* strains have a single protein annotated as FlaC<br>The FlaC of *C. rectus* is highly conserved, and differs from the FlaC of *C. showae*<br>*C. massilensis* has FlaC | Most *C. showae* strains have a single FlaC that aligns closely with *C. rectus* FlaC. | To be discussed further in C.H. Lago et al., in preparation) |
| **FlaC(s2)** |  | A variant form of FlaC only found in a few *C. showae* |  |
| **FlaG** | All *C. rectus* strains have a single protein annotated as FlaG | All *C. showae* strains have a single protein annotated as FlaG<br>*C. massilensis* has FlaG |  |
| **T6SS Type VI Secretion System** | Cr all complete genomes(Cr1-Cr 11), Cr12, Cr16-Cr17 partial T6SS , Cs7 and Cs17 (partial)<br>No T6SS found in Cm1<br>See Figure 5 to see the structure of the T6SS cluster visually. | At least three groups of T6SS in *C. rectus*. Cr1 matches best with Cs7, all other *C. rectus* in in either group 2 or 3 based on orientation of the clusters. | Most *C. showae* lack T6SS, the Cs7-CAM T6SS matches most closely with Cr1-ATCC 33238/RM3267. There are no repeats flanking T6SS in these two strains while all the other strains have repeats flanking the T6SS region and many have a phage lysin gene adjacent to the repeats. See supplemental Figure for details |
| **tssL/VasF (1)**<br>**tssK/VasE/ImpJ (2)**<br>**tssJ/VasD (3)** |  | These three genes are transcribed in the upstream direction away from the rest of the cluster.<br>TssL Membrane, Complex, TssK: Baseplate, TssJ Membrane Complex | The gene order for the *C. rectus* T6SS cluster does not match that reported for *C. jejuni* T6SS. In the legend for this table the order of the T6SS genes of *C. jejuni* is shown to indicate the differences. In addition to gene order, the hcp and VgrG genes are close to the T6SS cluster in *C. jejuni* and not separated from the rest of the cluster. |
| **tssA/ImpA/VasJ (4)**<br>**tssB/ImpB/VipA (5)**<br>**tssC/ImpC/VipB (6)** | Tss A docks to membrane complex, recruits baseplate complex | These genes and all that follow are transcribed in the downstream direction.<br>TssB contractile sheath, TssC contractile sheath | The importance of the separation of Hcp and VgrG from the rest of the T6SS cluster in *C. rectus* will be examined in future studies of the hcp mutant strains as well as other wt strains with different lengths of separation. |
| **tssE/VCA0109 (7)**<br>**tssF/ImpG/VasA (8)**<br>**tssG/VasH (9)** |  | TssE, TssF, and TssG Baseplate | VCA0109 is from *Vibrio cholerae* T6SS |
| **TssM/lcmF related (10)**<br>**TagH(11)** | Found in all complete T6SS clusters, TssM nomenclature only used for Cr1 and Cs7 all other Cr strains use lcmF related. | Verified all TssM match with lcmF-related by AA sequence alignment (data not shown) | TssM: Membrane Complex<br>TagH: Tip of Expelled Spike<br>TagH is annotated as TagA in *C. rectus* 33238, but we have chosen TagH since that is more common for T6SS. |
| **Hcp/TssD (12)**<br>**Hemolysin Co-regulated protein**<br>**VgrG/TssI\* (13)** | Both secreted effector and part of the secretion system apparatus<br>Secreted effector | Hcp and VgrG are adjacent to each other but separated from the rest of the cluster by 10,700-14,930 basepairs | *in 33238 VgrG was annotated as TssI |
| In comparison the T6SS of *C. jejuni* has the gene order of # 11, #10, #12, #1, #2, #3, #4, #5, #6, #7, #8, #9, #13 as defined in this table.<br>There is no gap between Hcp (12) or VgrG (13) and the rest of the cluster. So this represents a dramatic difference in the T6SS of *C. rectus* and that of *C. jejuni*. Of interest all hcp genes in *C. rectus* and *C. showae* are adjacent to VgrG unlike the separation seen in the *C. jejuni* arrangement. |  |  |  |

The protein or proteins that are secreted via the second type I system are suggested to be bacteriocins, so this might help in niche competition. It is of interest that *C. rectus* has two Type I systems while *C. showae* only have one per strain, i.e., either S-layer related or not related to S-layer secretion. These two systems are separated in the genome. *C. massilensis* the newest oral *Campylobacter* strain, has both types of T1SS and thus resembles *C rectus* more closely for these secretion systems. Indeed the crs gene is found in *C. massilensis* rather than the organization of multiple S-layer genes that are found in C. showae strains.

Type 2 protein secretion has not been reported in *C. rectus* and only one T2SS protein was identified in the isolates: PulG, a secretion envelope pseudopilin protein. However, various Type IV Pilus System (T4PS) genes were identified including PilB, PilC, PilO, and PilQ. MshM, a T4PS retraction ATPase was also identified. The T2SS, Type 4 pilus system (T4PS), archaeal flagella, and transformation system are evolutionarily related (Peabody et al., 2003). The T2SS/T4PS has been suggested to be involved with natural transformation in *C. jejuni* (Wiesner, Hendrixson, & DiRita, 2003) *C. rectus* 314 has been described to demonstrate this process (Wang, 2000). These two genome regions are shown in Supplemental Figure 3. The Type II/Type IV fimbrial assembly gene clusters were found in many *C. rectus* and *C. showae*, but the presence of S-layer in many of those same strains may limit natural transformation as a means of DNA uptake. Whether the presence of a T4SS together with a T2SS/T4PS aids in DNA uptake in *C. rectus/C. showae* is an interesting avenue for future research.

*C. rectus* and other *Campylobacter* species, including *C. showae and C. jejuni* have been shown to have a flagellar T3SS as compared to an injectosome-like T3SS like *Yersinia pestis* (Kreling, Falcone, Kehrenberg, & Hensel, 2020). The flagellum consists of basal body, hook, and filament.

*C. jejuni* has a class of virulence factors secreted through a flagellar T3SS, *Campylobacter* invasion (Cia) proteins. There are eight known Cia proteins in *C. jejuni* (CiaA-H) which have been shown to be required for full invasion of host cells by some *C. jejuni* strains. *C. rectus* only has one identified Cia protein: *CiaB*. A search among the 14 *C. showae* isolates showed that it has the *CiaB* protein as well. The role *CiaB* plays in adherence and invasion of *njc*-*Campylobacters* is unknown. Further work will be done to determine the secretion status of *CiaB* to determine if the antigen is secreted through the flagella similar to *C. jejuni* (Hughes Lago et al., Manuscript in Preparation). Table 5 lists the proteins that might be secreted through a *C rectus/C showae* flagellar T3SS since they have been shown to exit *C. jejuni* and related enteriic *C*ampylobacters via this process.The addition of pseudaminic acid to flagellin has not been previously addressed for *C. showae* and *C. rectus*, but all strains with flagellin genes have pseudaminic acid clusters as well, and also the MAF (Motility Acessory Protein) which has been shown to be the glycosyl transferase for Pse to flagellin.

In *Campylobacter* species, the T4SS is typically associated with horizontal gene transfer (HGT), which is important for genomic diversity (Wilson et al., 2009). Mutations in *C. jejuni* VirD4, a hallmark T4SS protein resulted in wild type natural transformation activity in *C. jejuni*, therefore the T4SS likely doesn’t play a major role for DNA uptake in this organism (Wiesner, Hendrixson, & DiRita, 2003)(Golz & Stingl, 2021). Instead, it appears to play a role in conjugation, a mechanism of HGT which occurs during cell-cell contact. *C. fetus venerealis* has a Type IVa class of T4SS which was shown to be involved in conjugation (Kienesberger et al., 2011). Of the *C. rectus* genomes analyzed, five isolates including SRR, OH, 314, 48803, and 27498 do not contain the hallmark genes required for a functional T4 system. These genes include VirB4, VirB9, VirB10, VirB11, VirD4, and IncQ/TraQ. In both *C. fetus* and *C. showae* lack of T4SS resulted in less virulence of these strains compared to strains containing T4SS (Gorkiewicz et al., 2010)(Hsu et al., 2019). This is something we are interested in testing functionally for *C. rectus*. We would expect for example, that T4SS+ *C. rectus* strains would be more invasive to host cells based on these previous studies. The two *C. showae* strains that we sequenced, CCUG 11641 and LMG 12636 both have T4SS genes. Indeed CCUG 11641 has two T4SS clusters, one termed “original Vir” and the other “duplicated Vir”. LMG 12636 has the duplicated Vir only. In the previous comparison of T4SS in *C showae*, three related strains 129_VTPP, 129_VTPPs, and 129_MSG demonstrated two Vir clusters, but the second set of these clusters were plasmid associated (Hsu et al., 2019). In our analysis, these duplicated clusters do not appear functional as they have overlapping repeats in some of the genes. Only the CCUG 11641 genome has two chromosomal T4SS clusters, and its duplicated cluster matches the single Vir cluster found in LMG 12636 (data not shown).

Results for the newer BV-BRC strain additions regarding Type IV are shown in Table 6. The percentage of strains exhibiting Type IV appears to be similar to that seen in other *Campylobacter* species, i.e., *C. jejuni*, and *C. fetus* subsp. venerealis (Panzenhagen, Portes, Dos Santos, Duque, & Conte Junior, 2021)(Silva et al., 2021). It should be noted that in *C. fetus* and *C. jejuni* the T4SS are often associated with plasmids.

**Table 6.** DNA Related Secretion System Clusters.

| Protein Name/<br>Locus name | Strains /No of strains |  | Unique features |
| --- | --- | --- | --- |
| **Type II Secretion** | In *C. jejuni* this system operates for natural transformation |  |  |
| **ComEA** | All *C. rectus* Isolates have 160 aa ComEA | Variable sizes of ComEA in *C. showae* |  |
| **PilG** | Some isolates have two proteins annotated as the T2S envelope pseudopilin protein within each strain. |  | Type II secretion envelope pseudopilin protein (PilG, guides folded protein to PilD in outer membrane) |
| **PilB/ctsE, PilC/ctsF** |  |  | Type IV fimbrial assembly, ATPase PilB<br>Type IV fimbrial assembly, ATPase PilC |
| **PilO, PilQ/ctsD** | pilO – putative periplasmic protein (required for glycosylation in *P. aeruginosa*) | mshL – Type IV pilus biogenesis protein<br>PilQ |  |
| **T4SS, Type IV Secretion System** | resembling Vir cluster from *Agrobacterium tumefaciens*<br>Now thought to be involved with DNA transfer through conjugation in *C. jejuni* |  | Has been suggested to be involved with protein secretion in *C. fetus* subsp. *veneralis* along with DNA transfer via conjugation |
| **T4SS Vir cluster-original** | *C. rectus* Cr1-Cr6; Cr, 11, Cr13, Cr18, Cr20<br>Cs, Cs3, Cs6a-c, Cs9, Cs12 | See below for original Vir genes | All Vir clusters in *C. rectus* strains appear to be in the same genomic location in all strains but Cr1 and Cr2 are in one group because the match falls off 5' to the Vir cluster while Cr3-Cr6 have close to 100kb of matching sequences (match on both sides of the Vir cluster) |
| **IncQ/TraQ (RP4 TrbM homolog) and Doc toxin (death on curing)** | One set of genes associated with original Vir cluster genes. All *C. rectus* with IncQ/TraQ and *C. showae* Cs7 (CAM) also have Doc | These two genes are closest to VirB4 and suggestive of the source of the original Vir cluster | Suggests a plasmid origin for these genes since both TraQ and Doc are associated with T4SS system plasmids transferred by conjugation |
| **VirB4, VirB6, VirB8** | *C. rectus* Cr1-Cr6<br>*C. showae* Cs1, Cs3- Cs6a-c, Cs9, Cs12 | ATPase required for both assembly of type IV secretion complex and secretion of T-DNA complex, VirB4 | VirB8 annotated as DNA competency protein for Cr1-Cr6 |
| **VirB9, VirB10, VirB11** |  | Outer membrane and periplasm component of type IV secretion of | Inner membrane protein of type IV secretion of T-DNA complex, TonB-like, VirB10 |
|  |  | T-DNA complex, has secretin-like domain, VirB9 | ATPase required for both assembly of type IV secretion complex and secretion of T-DNA complex, VirB11 |
| **VirD4** |  |  | Coupling protein VirD4, ATPase required for T-DNA transfer |
| **EcoRI modification methylase (EcoRI<sup>mm</sup>), Plasmid partitioning protein A (ParA), single stranded DNA binding protein (ssb), DNA topoisomerase (topo)** | Cr1-Cr6, Cs1, Cs3, Cs6a-c, Cs12 |  | Further suggests a plasmid origin for these genes since they relate to modification of plasmid DNA (EcoRI<sup>mm</sup>), subsequent intergration events (ssb, topo), and transfer to daughter cells (ParA) |
| **T4SS Vir cluster-duplicate** | Cs2-Cs3, Cs6a-6c |  | Cs2 has <u>only</u> the duplicate cluster and lacks the original cluster, the Cs2 Vir cluster most resembles the duplicated cluster from Cs3, the Cs6a-c duplicates are present on plasmids. |

The Type VI secretion system (T6SS), is a contact dependent nanomachine that resembles an inverted bacteriophage tail capable of secreting effector proteins to both prokaryotic and eukaryotic cells depending on bacterial species. First identified in *Vibrio cholerae* and *Pseudomonas aeruginosa*, it has since been found to be present in 25% of gram-negative bacterial species (Bingle, Bailey, & Pallen, 2008). No research has been done to elucidate the function(s) of T6SS in *C. rectus*. T6SS proteins identified in our panel of isolates include hcp (TssD) and VgrG (TssI), which are considered hallmark proteins of the T6SS. Each isolate also contains TssA (VasJ/ImpA), TssB (VipA/ImpB), TssC (VipB/ImpC), TssF (VasA/ImpG), TssG (VasB/ImpH), TssJ (VasD/Lip/SciN), and TssL (IcmH/DotU/VasF). All isolates also have an uncharacterized protein similar to VCA0109, which is required for hcp secretion in *V. cholerae* (Zheng, Ho, & Mekalanos, 2011). Further research must be done to test for functionality of the T6SS system in *C. rectus* and to determine how it works and whether it is anti-eukaryotic or anti-bacterial. The results for T6SS of all *C. rectus* strains in BV-BRC are shown in Table 5.

With the exception of *C. showae* CAM, none of the *C. showae* isolates have all the T6SS proteins. *C. showae* CAM has all the proteins for T6SS and a genomic comparison of the region shows very similar location of this cluster to *C. rectus* strains ATCC 33238 and OH2158. The lack of a complete T6SS in all but one *C. showae* strain suggests that the acquisition of this system occurred after the separation of the two species. CAM is notably more similar in other regions to some *C. rectus* strains. In particular, for the T6SS sequences, CAM is most similar to ATCC 33238, and this appears to be true for the only other *C. showae* strain presenting some of the T6SS genes including hcp, VgrG, IcmF (see Table 5). The presence of T6SS in at least two strains of *C. showae* may indicate that the percentage of strains with T6SS will increase as more showae genomes are added since not all *C. jejuni* strains have T6SS and the situation is more like the T4SS gene clusters. The *C. rectus* situation is therefore of special interest since even though a smaller number of genomes are available, a higher percentage of T6SS genes and full clusters have been identified overall than for *C. showae*. The location for T6SS is different for some strains of *C. rectus*, at least as it relates to one arm of the 30Kb region containing the cluster. In the CCUG strains 11640-11645, all isolates have one matching arm that ends in a conserved phage lysin resembling muramidase from phage λ. The ATCC 33238 group (3 strains) also exhibits a phage lysin but there is substantial deviation in the sequence of this enzyme, so it is unclear what the source is (Supplemental Figure 4).

## 4 CONCLUSION

Based on our comparative genome studies, we can see a clear demarcation between *C. rectus* and *C. showae* genomes. In particular, the TISS system for surface layer is present uniformly in complete *C. rectus* genomes while in complete *C. showae* genomes this system is only found in a subset of strains suggested to be more pathogenic. Additionally, the second T1SS system is uniformly distributed in *C. rectus*, but only found in *C. showae* strains that lack the S-layer type I system. This could be the result of divergent evolution or related to the high CRISPR content that has resulted in genome rearrangements in these two related species. There has been a suggestion that some of this diversity is due to the body location the strains were isolated from (i.e. oral vs. GI), but with the addition of the newest strains in BV-BRC this will have to be examined more closely. *C. massilensis* usually is more closely related to one group of *C. showae* strains (with CCUG 11641 as a member), and there is a clear separation of the two groups. So, the finding of two TISS and also the crs S-layer, in *C. massilensis* ssuggests another story for this newly discovered species.

T4SS is another interesting region for *C. rectus* versus *C. showae*. Several strains of *C. showae* demonstrated two T4SS regions. None of the *C. rectus* strains had two complete copies of T4SS, and one of the *C. showae* stains we had sequenced has two copies but only one of these copies was found in the other *C. showae* strain.

Finally, the T6SS appears to be largely *C. rectus* specific, in that only one completeT6SS cluster sequence was found in a *C. showae* strain, CAM. This *C. showae* strain often tracked more closely in other genomic regions to *C. rectus*, so perhaps was once a *rectus_showae* hybrid, or is an intermediate strain between *C. rectus* and *C. showae* evolutionarily. Although there are only a small number of *C. rectus* genomes available for study, it is striking that even incomplete genomes contain T6SS genes with IcmF being the most prominent one (Table 5). If this trend continues with the addition of more complete *C. rectus* genomes, the situation will resemble that of *V. cholerae* where most strains whether associated with cholera or not demonstrate T6SS clusters. *V. cholerae* utilizes Type VI for niche advantage as well as for host pathogenicity. With the 33238 *hcp* deletion mutant, we can investigate whether there are immunity groups in *C. rectus* for T6SS. This mutant can also support assessment of the importance of T6SS on host response including cell death.

Although the initial interest in genomic comparisons for *C. rectus* were aimed at investigating virulence factors and pathogenic pathways, a focus on secretion system gene clusters, has helped to demonstrate some unique features in the closely related oral strains of *C. rectus* and *C. showae.* It also allowed us to place *C. massilensis* as perhaps an intermediate between *C. showae* and *C. rectus*, rather than more closely related to *C. showae* as had been suggested during this organism’s discovery. Both *C. rectus* and *C. showae* have groups of strains that are more closely related to each other than to the whole group and this allows a selection from strains that belong to specific groups to examine phenotypes related to pathogenicity. Additionally, there are now additional classification schemes available for the *C. showae* and *C. rectus* strains based on their secretion systems and organization of those in different strains. There are now sufficient numbers of strains from both *C. rectus* and *C. showae* in BV-BRC to allow for powerful inter-strain comparisons to determine what genomic features are needed for full virulence. Additionally, the availability of 40+ genomes for the *rectus* and *showae* groups, will allow deeper comparisons with other *Campylobacter* species of importance to human health, *C. jejuni* and the two *C. fetus subsp.* for example.

## ACKNOWLEDGEMENTS

Funding information: NIH/NIDCR grant number R03DE023882. The Funding agency had no role in study design, data collection and interpretation, or the decision to submit the work for publication. No conflicts of interest to report for any of the authors of this manuscript. Dr. Drew Hillhouse of the TIGSS Genomics Core Facility generated the genome sequences. The first author, Casey Hughes Lago has completed fundamental bioinformatic research contributing to this manuscript, she created many of the figures presented such as BRIG and phylogenetic trees. She also developed assays and methods for working with *Campylobacter rectus* strains for biological assays (adherence and invasion, biofilm, pulsed field gel analysis). This project is part of Ms. Hughes Lago’s PhD studies. Dr Deborah Threadgill directed much of the bioinformatic studies and also conducted those of special interest based on her many years of working with *Campylobacter* species. She assisted Casey with figure generation as regards showing the gene order and conserved relationships across strains, and trying to understand the origin of the secretion systems. Both Ms. Hughes Lago and Dr. Threadgill worked on writing the manuscript, preparing tables and figures and making sure that all data was checked and re-checked for accuracy. Dr. Dana Blackburn was involved with the intial preparation of the *C. rectus* /*C. showae* genomic DNAs for sequencing and used assembly softoware for scaffolding prior to deposition of genome sequences in the RAST server, she also developed the RT-PCR for *C. rectus* and used this for mutant validation (ciaB). Ms. Kinder-Pavlicek helped with the RT-PCR assay development and studied some of the earliest genome information regarding the T6SS clusters. She also validated the hcp mutant and performed some experiments with it in completion of her MS degree.

We thank Ms. Kayla Beirne and Ms. Adelaide Sorbo for general lab support. For contributions from North Carolina State University, we thank Ms. Erin Harrell, Dr. Bridget Conley (Former NCSU undergraduate student, now: Assistant Professor at River Valley Community College) and Dr. Kristen Delaney-Ngyuen (now: Associate Professor at Fayetteville State University) for their involvement in mutant strain creation.

## REFERENCES

Alikhan, N., Petty, N. K., Ben Zakour, N. L., & Beatson, S. A. (2011). BLAST ring image generator (BRIG): Simple prokaryote genome comparisons. BMC Genomics, 12(1), 402. doi:10.1186/1471-2164-12-402

Anne. C. R. Tanner, Badger, S., Lai, C. H., Listgarten, M. A., Visconti, R. A., & Socransky, S. S. (1981). Wolinella gen. nov., wolinella succinogenes (vibrio succinogenes wolin et al.) comb. nov., and description of bacteroides gracilis sp. nov., wolinella recta sp. nov., campylobacter concisus sp. nov., and eikenella corrodens from humans with periodontal disease. International Journal of Systematic Bacteriology, 31(4), 432–445.

Antezack, A., Boxberger, M., Rolland, C., Ben Khedher, M., Monnet-Corti, V., & La Scola, B. (2021). Isolation and characterization of campylobacter massiliensis sp. nov., a novel campylobacter species detected in a gingivitis subject. International Journal of Systematic and Evolutionary Microbiology, 71(10) doi:10.1099/ijsem.0.005039

Arndt, D., Grant, J. R., Marcu, A., Sajed, T., Pon, A., Liang, Y., & Wishart, D. S. (2016). PHASTER: A better, faster version of the PHAST phage search tool. Nucleic Acids Research, 44(W1), W16–W21. doi:10.1093/nar/gkw387

Bacon, D. J., Alm, R. A., Burr, D. H., Hu, L., Kopecko, D. J., Ewing, C. P., … Guerry, P. (2000). Involvement of a plasmid in virulence of campylobacter jejuni 81-176. Infection and Immunity, 68(8)

Bertelli, C., Laird, M. R., Williams, K. P., Lau, B. Y., Hoad, G., Winsor, G. L., & Brinkman, F. S. L. (2017). IslandViewer 4: Expanded prediction of genomic islands for larger-scale datasets. Nucleic Acids Research, 45(W1), W30–W35. doi:10.1093/nar/gkx343

Bingle, L. E., Bailey, C. M., & Pallen, M. J. (2008). Type VI secretion: A beginner’s guide. Current Opinion in Microbiology, 11(1), 3–8. doi:10.1016/j.mib.2008.01.006

Bleumink-Pluym, N., Van Alphen, L. B., Bouwman, L. I., Wösten, M. M., & Van Putten, J. P. M. (2013). Identification of a functional type VI secretion system in campylobacter jejuni conferring capsule polysaccharide sensitive cytotoxicity. PLoS Pathogens, 9(5), e1003393. doi:10.1371/journal.ppat.1003393

Bobetsis Yiorgos, A., Barros, S. P., & Offenbacher, S. (2006). Exploring the relationship between periodontal disease and pregnancy complications. The Journal of the American Dental Association, *137*

Brettin, T., Davis, J. J., Disz, T., Edwards, R. A., Gerdes, S., Olsen, G. J., … Xia, F. (2015). RASTtk: A modular and extensible implementation of the RAST algorithm for building custom annotation pipelines and annotating batches of genomes. Scientific Reports, 5(1), 8365. doi:10.1038/srep08365

Castaño-Rodríguez, N., Kaakoush, N. O., Lee, W. S., & Mitchell, H. M. (2017). Dual role of helicobacter and campylobacter species in IBD: A systematic review and meta-analysis. Gut, 66(2), 235–249. doi:10.1136/gutjnl-2015-310545

Costa, D., & Iraola, G. (2019). *Pathogenomics of emerging campylobacter species* American Society for Microbiology. doi:10.1128/cmr.00072-18

Doub, J. B. (2021). A rare case of campylobacter rectus pyogenic extensor tenosynovitis. Germs (Bucureşti), 11(4), 604–607. doi:10.18683/germs.2021.1296

Golz, J. C., & Stingl, K. (2021). Natural competence and horizontal gene transfer in campylobacter. Fighting campylobacter infections (pp. 265–292). Cham: Springer International Publishing. doi:10.1007/978-3-030-65481-8_10 Retrieved from https://library.biblioboard.com/viewer/03a14878-b782-11eb-b971-0a9b31268bf5

Gorkiewicz, G., Kienesberger, S., Schober, C., Scheicher, S. R., Gully, C., Zechner, R., & Zechner, E. L. (2010). A genomic island defines subspecies-specific virulence features of the host-adapted pathogen campylobacter fetus subsp. venerealis. Journal of Bacteriology, 192(2), 502–517. doi:10.1128/JB.00803-09

Hodges, F. J., Torres, V. V. L., Cunningham, A. F., Henderson, I. R., & Icke, C. (2023). Redefining the bacterial type I protein secretion system. Advances in Microbial Physiology, doi:10.1016/bs.ampbs.2022.10.003

Hsu, T., Gemmell, M. R., Franzosa, E. A., Berry, S., Mukhopadhya, I., Hansen, R., … Hold, G. L. (2019). Comparative genomics and genome biology of campylobacter showae. Emerging Microbes & Infections, 8(1) doi:10.1080/22221751.2019.1622455

Hughes Lago, C., Threadgill, D.S., (Manuscript in Preparation). Investigating the role of Campylobacter rectus CiaB in adherence, invasion, and host response.

Hung, D. L. L., Teng, J. L. L., Lau, S. K. P., & Woo, P. C. Y. (2019). Lemierre’s syndrome associated with campylobacter rectus bacteremia. Infectious Microbes and Diseases, 1(1), 27–29. doi:10.1097/IM9.0000000000000004

Kaakoush, N. O., Castaño-Rodríguez, N., Man, S. M., & Mitchell, H. M. (2015). Is campylobacter to esophageal adenocarcinoma as helicobacter is to gastric adenocarcinoma? Trends in Microbiology (Regular Ed.), 23(8), 455–462. doi:10.1016/j.tim.2015.03.009

Kato Yukio, Shirae Mitsuyuki, Murakami Masaru, Mizausawa Tetsuya, Hagimoto Atsuki, Wada Koichiro, … Asai Furmitoshi. (2011). Molecular detection of human periodontal pathogens in oral swab specimens from dogs in Japan. J Vet Dent, 28(2), 84–89.

Kienesberger, S., Schober Trummler, C., Fauster, A., Lang, S., Sprenger, H., Gorkiewicz, G., & Zechner, E. L. (2011). Interbacterial macromolecular transfer by the *Campylobacter fetus* subsp. venerealis type IV secretion system. Journal of Bacteriology, 193(3), 744–758. doi:10.1128/JB.00798-10

Kreling, V., Falcone, F. H., Kehrenberg, C., & Hensel, A. (2020). *Campylobacter* sp.: Pathogenicity factors and prevention methods—new molecular targets for innovative antivirulence drugs? Applied Microbiology and Biotechnology, 104(24), 10409–10436. doi:10.1007/s00253-020-10974-5

Lam, J. Y. W., Wu, A. K. L., Ngai, D. C., Teng, J. L. L., Wong, E. S. Y., Lau, S. K. P., … Woo, P. C. Y. (2011). Three cases of severe invasive infections caused by *Campylobacter rectus* and first report of fatal *C. rectus* infection. Journal of Clinical Microbiology, 49(4), 1687–1691. doi:10.1128/JCM.02487-10

Lee, I., Ouk Kim, Y., Park, S., & Chun, J. (2016). *OrthoANI: An improved algorithm and software for calculating average nucleotide identity* Microbiology Society. doi:10.1099/ijsem.0.000760

Lertpiriyapong, K., Gamazon, E. R., Feng, Y., Park, D. S., Pang, J., Botka, G., … Fox, J. G. (2012). *Campylobacter jejuni* type VI secretion system: Roles in adaptation to deoxycholic acid, host cell adherence, invasion, and in vivo colonization. PloS One, 7(8), e42842. doi:10.1371/journal.pone.0042842

Macuchi, P. J., & Tanner, A. C. R. (1999). *Campylobacter* species in health, gingivitis, and periodontitis. J Dent Res, (79), 785–792.

O’Toole, G. A. (2011). Microtiter dish biofilm formation assay. Journal of Visualized Experiments, (47) doi:10.3791/2437

Panzenhagen, P., Portes, A. B., Dos Santos, A. M. P., Duque, S. D. S., & Conte Junior, C. A. (2021). The distribution of *Campylobacter jejuni* virulence genes in genomes worldwide derived from the NCBI pathogen detection database. Genes, 12(10) doi:10.3390/genes12101538

Peabody, C. R., Chung, Y. J., Yen, M., Vidal-Ingigliardi, D., Pugsley, A. P., & Saier, M. H. (2003). Type II protein secretion and its relationship to bacterial type IV pili and archaeal flagella. Microbiology Society. doi:10.1099/mic.0.26364-0

Rambout, A. (2007). FigTree, a graphical viewer of phylogenetic trees. Retrieved from: http://tree.bio.ed.ac.uk/software/figtree/

Rocha, D. J. P., Santos, C. S., & Pacheco, L. G. C. (2015). Bacterial reference genes for gene expression studies by RT-qPCR: Survey and analysis. Antonie Van Leeuwenhoek, 108(3), 685–693. doi:10.1007/s10482-015-0524-1

Salah Ud-Din, A. I. M., & Roujeinikova, A. (2017). Flagellin glycosylation with pseudaminic acid in *Campylobacter* and *Helicobacter*: Prospects for development of novel therapeutics. Cellular and Molecular Life Sciences, 75(7), 1163. doi:10.1007/s00018-017-2696-5

Shiga, Y., Hosomi, N., Nezu, T., Nishi, H., Aoki, S., Nakamori, M., … Fletcher, H. M. (2020). Association between periodontal disease due to *Campylobacter rectus* and cerebral microbleeds in acute stroke patients.(research article). PloS One, 15(10), e0239773. doi:10.1371/journal.pone.0239773

Silva, M. F., Pereira, A. L., Fraqueza, M. J., Pereira, G., Mateus, L., Lopes-Da-Costa, L., & Silva, E. (2021). Genomic and phenotypic characterization of *Campylobacter fetus* subsp. venerealis strains. Microorganisms, 9(2) doi:10.3390/microorganisms9020340

Spitz, O., Erenburg, I. N., Beer, T., Kanonenberg, K., Barry Holland, I., Schmitt, L., … Christie, P. J. (2019). Type I secretion systems-one mechanism for all? doi:10.1128/microbiolspec.PSIB-0003-2018.

Tu, Z., Gaudreau, C., & Blaser, M. J. Mechanisms underlying Campylobacter fetus pathogenesis in humans: Surface-layer protein variation in relapsing infections

Veyrine, A., Sturque, J., Zlowodski, A., Sourimant, T., Le Gal, M., & Denis, F. (2019). Pelvic peritonitis caused by *Campylobacter rectus* infection: Case report and literature review. Journal of Medical Cases, 10(4), 97–100. doi:10.14740/jmc3275

Wang, B. (2000). Molecular genetic analysis of the role of the s-layer from Campylobacter rectus in pathogenesis (PhD).

Warren, R. L., Freeman, D. J., Pleasance, S., Watson, P., Moore, R. A., Cochrane, K., … Holt, R. A. (2013). Co-occurrence of anaerobic bacteria in colorectal carcinomas. Microbiome, 1(1), 16. doi:10.1186/2049-2618-1-16

Wattam, A. R., Davis, J. J., Assaf, R., Boisvert, S., Brettin, T., Bun, C., … Stevens, R. L. (2017). *Improvements to PATRIC, the all-bacterial bioinformatics database and analysis resource center* Oxford University Press (OUP). doi:10.1093/nar/gkw1017

Wiesner, R. S., Hendrixson, D. R., & DiRita, V. J. (2003). Natural transformation of *Campylobacter jejuni* requires components of a type II secretion system. Journal of Bacteriology, 185(18), 5408–5418. doi:10.1128/JB.185.18.5408-5418.2003

Wilson, D. J., Gabriel, E., Leatherbarrow, A. J. H., Cheesbrough, J., Gee, S., Bolton, E., … Fearnhead, P. (2009). Rapid evolution and the importance of recombination to the gastroenteric pathogen campylobacter jejuni. Molecular Biology and Evolution, 26(2), 385–397. doi:10.1093/molbev/msn264

Yeo Alvin, Smith, M. A., Lin, D., Riche, E. L., Moore, A., Elter, J., & Offenbacher, S. (2005). Campylobacter rectus mediates growth restriction in pregnant mice. Journal of Periodontology, *76*

Zheng, J., Ho, B., & Mekalanos, J. J. (2011). Genetic analysis of anti-amoebae and anti-bacterial activities of the type VI secretion system in *Vibrio cholerae*. PLoS ONE, 6(8), e23876. doi:10.1371/journal.pone.0023876

