## Supplemental Information for "Comparative Genomic Analysis of *Campylobacter rectus* and Closely Related Species"

**RNA Isolation Protocol**. To isolate RNA, the following conditions were used: five mL of either MFF broth or tryptic soy broth + formate/fumarate (TSBFF) (tryptic soy broth 30 g/L, 0.2% sodium formate, 0.2% ammonium fumarate) was inoculated with each bacterial strain and cultures were grown anaerobically for 8 h at 37°C. Following incubation, the cultures were used to inoculate 50 mL of MFF or TSBFF broth to an $OD_{600}$=0.02. Cultures started at $OD_{600}$=0.02 were grown anaerobically overnight at 37°C until an $OD_{600}$=0.25 was reached; 3 mL from each culture was centrifuged at 14,800 rpm for 1 min, supernatant removed, and pellets resuspended in 750 μL of RNAlater (Thermo Fisher Scientific, Waltham, MA, USA). RNA was extracted with the RNeasy Mini Kit (Qiagen, Valencia, CA, USA) using manufacturer's instructions (Qiagen RNAprotect Bacteria Reagent Handbook) with some modifications. To eliminate any residual DNA, another DNase digest following extraction was performed using the DNA-free DNA Removal Kit (Thermo Fisher Scientific). 3 μL each of DNase and 10X DNase I Buffer was added to 24 μL of RNA from each sample and incubated for 90 min. Samples were centrifuged in RNAlater at 14,800 rpm for 1 min. The supernatant was removed and samples were resuspended in 200 μL of ice-cold lysis buffer (20mM Tris pH 7.5, 10mM EDTA pH 8.0, 150 mM β-mercaptoethanol, 0.5% SDS, 150 μg/mL proteinase K). After a 5 min incubation on ice, samples were incubated at 65°C for another 5 min. After lysis, steps 9-10 from protocol 4 were performed followed by protocol 7 (Qiagen RNAprotect Bacteria) including treatment with the RNase-Free

DNase Set (Qiagen) for on-column DNase digestion. An additional DNase digest was performed as described in the methods section of the main manuscript.

To test for the presence of DNA, PCR was performed using primers for *glyA* and *rpoA* targets (Insert Primer Table). The Qiagen Taq PCR core kit described above was used. The following parameters were used on a Bio-Rad C1000 Touch Thermal Cycler (Bio-Rad Laboratories, Hercules, CA, USA): 94°C for 3 min, then 35 cycles of 94°C 30 s, 55°C 30 s, 72°C 1 min, with a final elongation of 72°C for 5 min.

**Supplemental Table 1.** Primers Used in this Study

| Gene | Forward Primer | Reverse Primer |
|---|---|---|
| **16S** | CGGTACCCAAGGAATAAGCA | TCCTTTACGCCCAGTGATTC |
| ***gyrA*** | AAAAACGGCATCGTAAAACG | GCTCGTCGTTCTCGTCTAGG |
| ***recA*** | TCGGACTTGACTTGGCTCTT | GCGATGATATGAAGCGTGAG |
| ***glyA*** | AGCGCATATACGCGAGAGAT | TACGACTAGACCGGCGATGT |
| ***rpoA*** | CACGCGAAGTTGCTACAAAG | CATCCATCAAAGCAAGCTCA |
| ***rpoD*** | CCAGGATGGCAAAGTCAAAT | CCGATGTTGCCTTCTTGAAT |

**Supplemental Table 2.** BestKeeper Data

| Strain | Gene | GM [Cq] | (min, max) [Cq] | SD [± Cq] | (min, max) [x-fold] | SD [± x-fold] |
|---|---|---|---|---|---|---|
| **33238** | ***gyrA*** | 16.41 | 15.88,17.1 | 0.37 | -1.4,1.57 | 1.27 |
| | ***recA*** | 18.29 | 17.87,19.23 | 0.33 | -1.28,1.77 | 1.24 |
| | ***glyA*** | 21.43 | 20.04,24.69 | 0.87 | -2.13,6.34 | 1.76 |
| | ***rpoA*** | 12.76 | 12.37,13.36 | 0.23 | -1.26,1.45 | 1.16 |
| | ***rpoD*** | 16.66 | 14.93,25.22 | 2.14 | -2.11,82.27 | 3.99 |
| **314** | ***gyrA*** | 16.67 | 15.44,17.44 | 0.46 | -2.2,1.67 | 1.35 |
| | ***recA*** | 18.6 | 17.86,19.31 | 0.41 | -1.56,1.54 | 1.3 |
| | ***glyA*** | 35.38 | 34.14,37.09 | 0.59 | -1.99,2.62 | 1.46 |
| | ***rpoA*** | 13.00 | 11.46,13.80 | 0.46 | -2.53,1.65 | 1.35 |
| | ***rpoD*** | 18.37 | 17.77,19.11 | 0.42 | -1.34,1.45 | 1.31 |

**Supplemental Table 3.** GeNorm Analysis

| | Stability Value | |
|---|---|---|
| **Gene** | **33238** | **314** |
| ***gyrA*** | 0.008 | 0.016 |
| ***recA*** | 0.003 | 0.016 |
| ***glyA*** | 0.027 | 0.029 |
| ***rpoA*** | 0.012 | 0.032 |
| ***rpoD*** | 0.065 | 0.031 |

**Supplemental Table 4.** Distribution of Crispr Repeats, Spacers, and Arrays in *C. rectus* Isolates

| ***C. rectus* Strain** | **Crispr Repeats** | **Crispr Spacers** | **Crispr Arrays** |
|---|---|---|---|
| **11645** | 177 | 172 | 5 |
| **11643** | 182 | 178 | 4 |
| **11642** | 170 | 165 | 5 |
| **11640** | 119 | 115 | 5 |
| **48803** | 153 | 149 | 4 |
| **33238** | 85 | 83 | 2 |
| **314** | 31 | 29 | 2 |
| **7615** | 85 | 83 | 2 |
| **27498** | 12 | 10 | 2 |
| **OH2158** | 62 | 59 | 3 |
| **SRR9217473-mag-bin.5** | 18 | 17 | 1 |

**DNA Uptake Protocol.** 15 mL MFF broth tubes were inoculated with each *C. rectus* strain and incubated overnight for pre-growth in anaerobic conditions. After pre-growth, 50 mL of MFF broth was inoculated and grown to $OD_{600}$ 0.20 for each strain. Cells were harvested by centrifugation and washed twice with 10 mL of electroporation buffer (EB) (15% glycerol, 0.272 M sucrose, 0.57 mM $KH_2PO_4$, 2.43 mM $K_2HPO_4$, pH 7). Pellets were resuspended in 15 mL electroporation buffer and then $6x10^8$ cells were removed and centrifuged before resuspending in 60 µL of EB. 600 ng of pDK619 plasmid was added to each strain before transferring to 1 mL of MFF broth in glass tubes. Tubes were incubated in anaerobic condition for 4 h before centrifuging. Pellets were resuspended in 200 µL of PBS and then plated on MFF agar plates containing 40 µg/mL spectinomycin. After 48 h, CFUs were counted.

**Supplemental Table 5.** DNA Uptake Data

| Isolate | DNA Uptake Visible |
|---|---|
| **Strain 314** | |
| **Strain 33238** | + |
| **Strain 48803** | |
| **Strain 27948** | |
| **Strain 11643** | |
| **Strain 11645** | + |
| **Strain 11642** | + |
| **Strain 11640** | |
| **Strain 7615** | + |

**Supplemental Table 6.** Mutant Table with Molecular Technique Used

| Mutant | Name | Molecular Technique Used |
|---|---|---|
| **CiaB** | *Campylobacter* Invasion Antigen B | Gateway Cloning |
| **VirB9** | Outer membrane and periplasm component of T4S of T-DNA complex | Gibson Assembly |
| **Hcp** | Hemolysin Co-Regulated protein | Gibson Assembly |

### Supplemental Figures

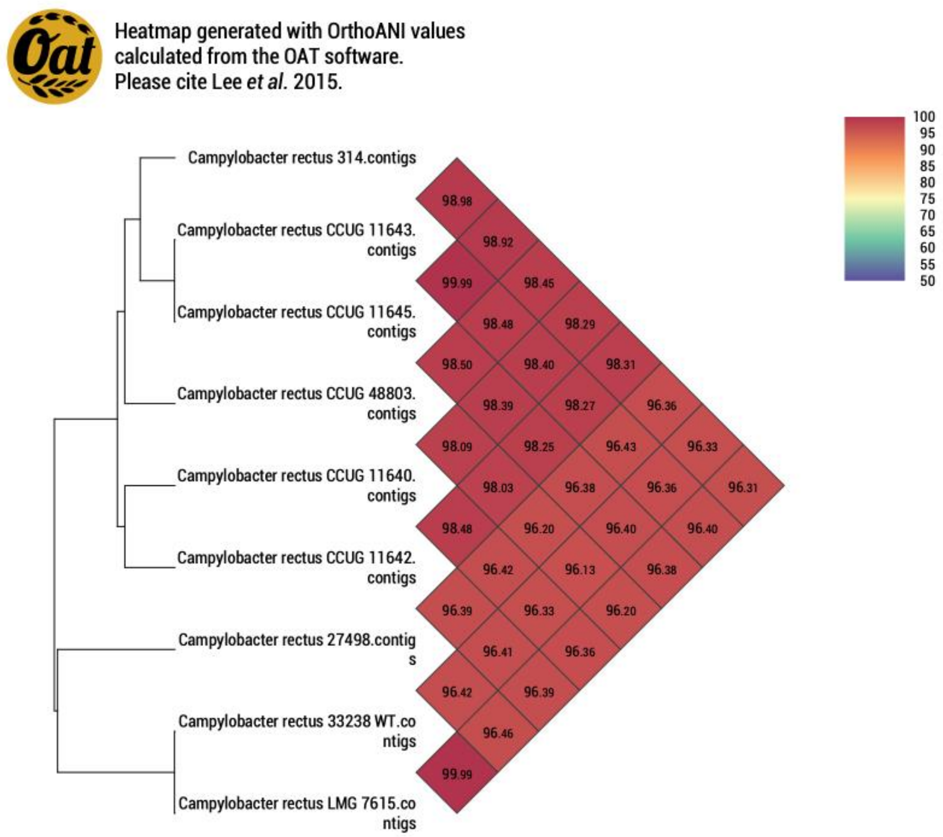


**Supplemental Figure 1.** Pairwise Comparisons between *C. rectus* Isolates

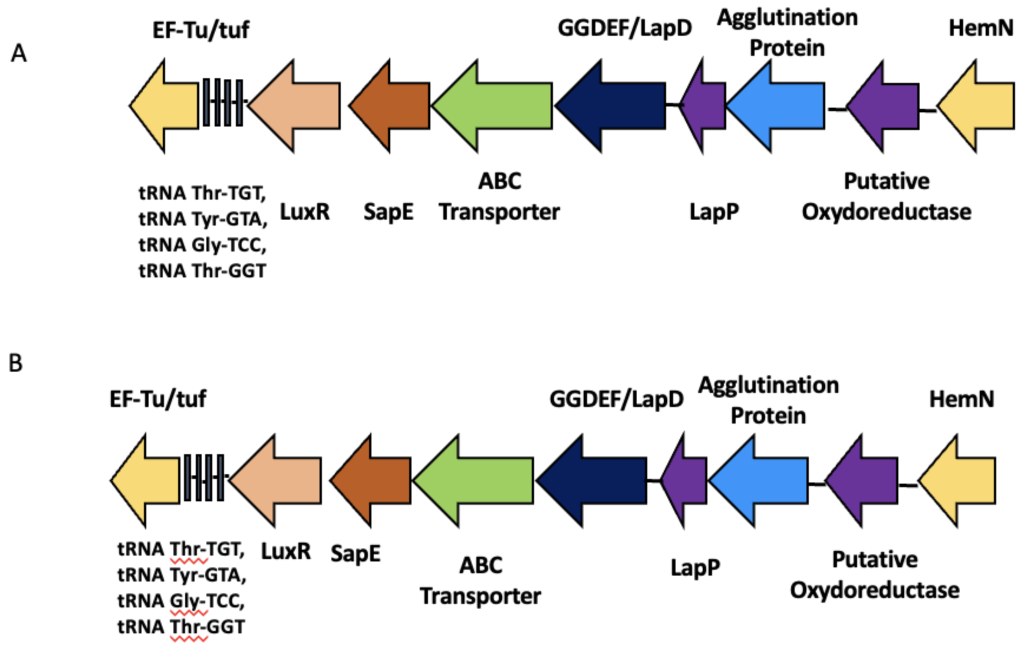


**Supplemental Figure 2.** Non-S-layer T1SS (possible bacteriocin secretion) A) Strain *C. rectus* 7615 and B) *C. massilensis*

(Gene Names/Abbreviations: SapE: T1S membrane fusion protein hylD family, EF-TU: translation elongation factor tu, LuxR: regulatory protein, ABC Transporter: transmembrane region:ABC transporter:Peptidase C39, bacteriocin processing, LapD/GGDEF: domain proteins, Putative oxidoreductase: ferredoxin-type protein, clusters with CPO, HemN: Coproporphyrinogen III oxidase, oxygen-independent, Hypo: hypothetical protein. *C showae* show very similar Type I Non-S-layer cluster organization to both *C. rectus* and *C. massilensis*)

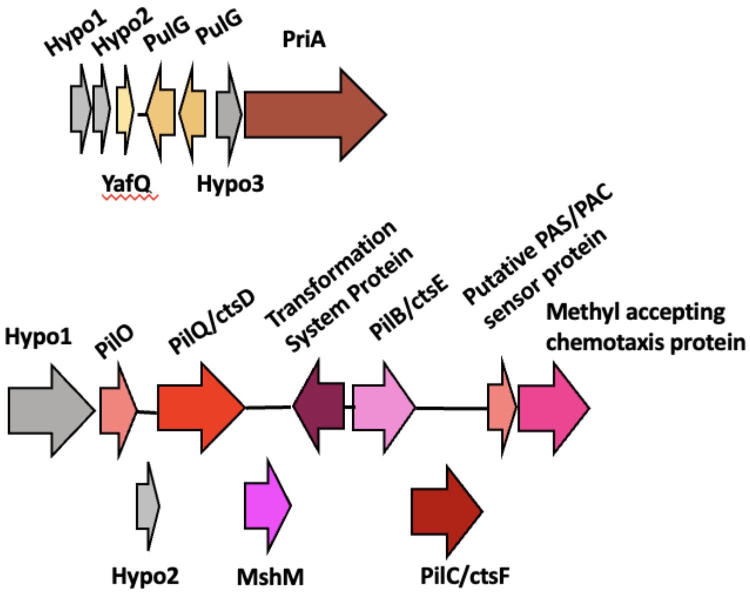


**Supplemental Figure 3.** T2SS Cluster in *C. rectus* 7615 (YafQ: Toxin Protein**,** PulG: Type II secretion envelope pseudopilin protein (PulG,guides folded protein to PulD in outer membrane)some strains have two of these**,** Helicase PriA: essential for OriC/DnaA independent DNA replication, B**.** Type IV Fimbrial assembly cluster pillO – putative periplasmic protein, PilQ: Type IV pilus biogenesis protein (mshL))

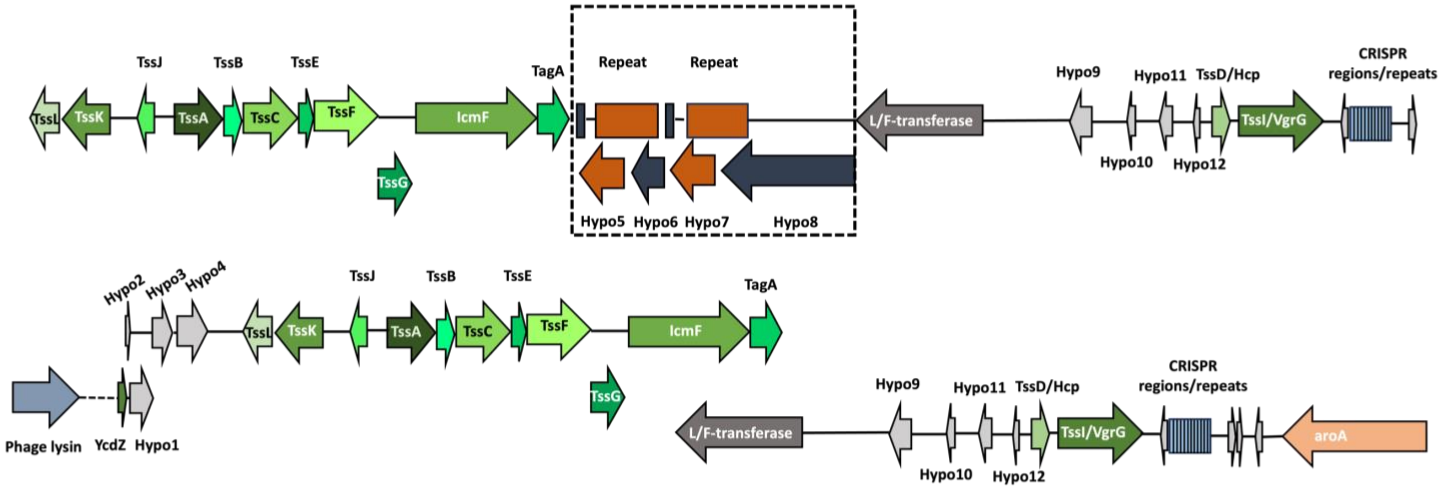


**Supplemental Figure 4.** Example of T6SS Gene Neighborhood in 11643 with the extended flanking regions (shown)

Gene Names and/or Orthologs: Phage lysin*; 1,4-beta-N-acetylmuramidase or lysozyme #Protein S in phage lambda, TssL; VasF/ImpK, TssK; VasE/ImpJ, TssJ; VasD, TssA; VasJ/ImpA, TssB; ImpB, TssC; VipB/ImpC, TssE; VCA0109, TssF; VasA/ImpG, TssG; VasB, TssM; VasK/IcmF, TagA, L/F-transferase; Leucyl/phenylalanyl-tRNA--protein transferase, aroA; chorismate synthase, Hypo; hypothetical protein.

*The phage lysin and YcdZ proteins are separated by 8 hypothetical proteins spanning 5623 base pairs.
